# Genetic Targeting and Conductance-Based Modeling Reveal Novel Diversity Within Mouse Type II Spiral Ganglion Neurons

**DOI:** 10.64898/2025.12.05.692182

**Authors:** Nathaniel Nowak, Radha Kalluri

## Abstract

Spiral ganglion neurons (SGNs) transmit auditory signals from the cochlea to the brain and are divided into two main types: type I and type II, distinguished by their anatomy and connectivity. However, the function of type II SGNs remains poorly understood due to their scarcity and lack of clear physiological markers. In this study, we use two Cre-dependent fluorescent reporter mouse lines to enhance the identification and targeting of type II SGNs for whole-cell patch-clamp recordings. We reveal a set of distinguishing biophysical features, most notably, the presence of an inactivating potassium current and weaker voltage-gated sodium currents, that clearly separate type II SGNs from their type I counterparts. Additionally, we uncover greater-than-expected heterogeneity among type II SGNs, including variation in size, excitability, and ion channel expression. These features suggest the existence of distinct subtypes of type II SGNs, with potential differences in function. We find that most type II SGNs are relatively unexcitable and incapable of repetitive firing. Instead, they appear to be better suited to integrating sustained signals, potentially supporting roles in detecting cochlear damage or modulating efferent feedback. Additionally, through computational modeling, we demonstrate that removing the inactivation component of the inactivating potassium current specific to type II SGNs allowed repetitive spiking to similar levels seen in type I SGNs, suggesting a crucial role for the current in stifling type II SGN activity. Together, our findings define biophysical signatures that distinguish SGN types and subtypes, offering new insight into their contributions to normal hearing and cochlear pathology.

**Significance:** The sensory neurons of the cochlea are divided into type I and type II spiral ganglion neurons. Type I spiral ganglion neurons convey the main features of sound information. The rarer type II spiral ganglion neurons appear to be putative auditory nociceptors, responding to cochlear damage. By combining genetic tools, electrical activity recordings, and computational models, we demonstrate that type I and type II spiral ganglion neurons have distinctive ion channel profiles and firing properties. Furthermore, we report previously undescribed ion channel diversity within the type II spiral ganglion neuron population, suggesting varied functions. Our results highlight the parallels between type II spiral ganglion neurons and somatosensory nociceptors and provide a framework for selectively targeting distinct auditory neuron populations.

## Introduction

Sensory systems translate environmental stimuli, such as chemical signals or physical energy, into neural activity interpretable by the central nervous system. In the mammalian auditory system, sensory hair cells transduce sound via basilar membrane motion in the cochlea (Fettiplace, 2017). This activates spiral ganglion neurons (SGNs), which relay information to the cochlear brainstem nuclei (Robertson, 1984; Brown, 1994; Nayagam et al., 2011).

The spiral ganglion comprises two primary neuron types distinguished by their cochlear hair cell targets. Type I SGNs have single unbranched dendrites that reliably generate action potentials in response to glutamate release from inner hair cells (IHCs) (Liberman, 1980; Rutherford et al., 2012; Wu et al., 2016). In contrast, type II SGNs extend long, branching dendrites to contact dozens of outer hair cells (OHCs) (Spoendlin, 1969; Berglund and Ryugo, 1987). Although they can be activated by OHC glutamate release, even synchronous input from all presynaptic OHCs rarely evokes an action potential (Weisz et al., 2012; Wood et al., 2020). Overall, type I and type II SGNs present two parallel pathways for distinct auditory information to flow from the cochlea to the brain. However, how type II SGNs process cochlear input and what specific role they play in auditory perception remain poorly understood.

Diversity within type I SGNs is now well established. Even neurons innervating the same IHC, thus encoding the same sound frequency, display anatomical (Liberman, 1982), transcriptional (Petitpré et al., 2018; Shrestha et al., 2018; Sun et al., 2018), and functional variability (Liberman, 1978; Siebald et al., 2023). Gradients in electrophysiological properties derived from variations in ion channel densities (Mo and Davis, 1997; Markowitz and Kalluri, 2020) are amongst the factors that influence the heterogeneity of firing patterns that subserve functional differences across so-called subtypes of type I SGNs.

By contrast, the molecular and biophysical properties of type II SGNs remain poorly understood. This is partly due to their relative scarcity and due to the technical difficulty of recording from their unmyelinated axons *in vivo* (Robertson, 1984; Brown, 1994) and dendrites *in vitro* (Weisz et al., 2009, 2012, 2014; Liu et al., 2015). Furthermore, the few recordings from stochastically sampled SGN somata suggest that type II SGNs are biophysically distinct from type I SGNs (Jagger and Housley, 2003; Reid et al., 2004; Markowitz and Kalluri, 2020). Additional evidence from electrophysiological recordings (Liu et al., 2015), c-Fos staining (Flores et al., 2015; Weisz et al., 2021), and calcium imaging (Nowak et al., 2021) implicates type II SGNs in sensing noxious or damaging stimuli within the cochlea. Collectively, these findings support the view that type II SGNs convey sensory information distinct from type I SGNs.

Recently, several Cre-driver lines have enabled selective labeling of type II SGNs using fluorescent reporters (Vyas et al., 2017, 2019; Wu et al., 2018). Here, we use two such lines to facilitate patch-clamp recordings from type II SGN somata. The first, *Serotonin Reuptake Transporter*-Cre (SERT^Cre^), drives fluorescent tdTomato expression selectively in type II SGNs, providing a visual means to distinguish them from type I SGNs. However, this line is not inclusive of all type II SGNs (Vyas et al., 2019). To improve coverage, we also use the *Tachykinin 1*-Cre (Tac1^Cre^) line (Nowak et al., 2021; Sanders and Kelley, 2022), which labels a wider population, which is expected to be more broadly inclusive of type II SGNs but potentially some type I SGNs.

This study leverages recordings from both fluorescent reporter lines to identify the key distinguishing biophysical properties between type I and type II SGNs. Our results confirm known distinctions between these cell types and further reveal previously underappreciated biophysical and morphological diversity within the type II SGN population, suggesting that, like type I SGNs, type II SGNs may comprise functionally heterogeneous subtypes.

## Methods

### Animal models

Cochlear epithelia, including the spiral ganglion, were collected from C57BL/6J mice of either sex between postnatal days 5 and 10 (P0 = day of birth). Breeding pairs consisted of homozygous *Serotonin Reuptake Transporter*-Cre (SERT^Cre^) mutant mice provided by Jackson Laboratory (RRID:IMSR_JAX:014554) or *Tachykinin1* (Tac1^Cre^) (RRID:IMSR_JAX: 021877) and homozygous Ai14 Cre-reporter mice (RRID:IMSR_JAX:007914). Some control animals were bred by pairing two wild type C57Bl6/J mice or by pairing an Ai14 mouse with a wild type C57/Bl6J mouse. All animals were housed and handled in accordance with the National Institutes of Health Guide for the Care and Use of Laboratory Animals, and all procedures were approved by the University of Southern California Institutional Animal Care and Use Committee.

### Viral injections

Intracerebroventricular viral injections were performed at P2 in one male and one female heterozygous Ai14 mouse using a previously described protocol (Kim et al., 2014). We injected pENN.AAV.hSyn.HI.eGFP-Cre.WPRE.SV40 (a gift from James M. Wilson, Addgene viral prep # 105540-PHPeB; http://n2t.net/addgene:105540; RRID:Addgene_105540), diluted 1:1000 to an estimated final titer of ∼1 × 10^10^ vg/mL. This dilution permitted sparse, stochastic Cre expression in spiral ganglion neurons, resulting in tdTomato expression independent of SERT^Cre^ or Tac1^Cre^ driver lines.

### Immunohistochemistry

Otic capsules were collected from euthanized SERT^Cre^; Ai14, Tac1^Cre^; Ai14, and virus-injected Ai14 mice between P17 and P87. Tissues were fixed in 4% paraformaldehyde for 1 hour at room temperature and rinsed three times with phosphate-buffered saline (PBS). Fixed otic capsules were decalcified in 10% EDTA for 16–24 hours, followed by additional PBS washes. Cochlear epithelia and attached spiral ganglia were then dissected out and cut into separate turns in PBS.

Free-floating cochlear turns were blocked in 2% Triton-X in goat serum dilution buffer (GSDB) for 1 hour at room temperature. Samples were incubated for 72 hours at 4°C with primary antibodies diluted in GSDB containing 0.3% Triton-X, followed by three 10-minute PBS washes. Secondary antibody incubation was performed overnight (16-24 hours) at 4°C, followed by three 10-minute PBS washes.

Primary antibodies were anti-Calretinin (rabbit polyclonal, Swant #7699/3H; 1:1,000) and anti-β-Tubulin III (mouse monoclonal IgG2a, BioLegend #801202; 1:200). Secondary antibodies, used at 1:200 dilution, were Alexa Fluor 488 anti-rabbit (Invitrogen A11008) and Alexa Fluor 647 anti-mouse IgG2a (Invitrogen A21241).

#### Imaging

Immunostained samples were mounted in Vectashield Antifade Mounting Medium (Vector Laboratories). Raised dots of clear nail polish were used to support coverslips and prevent compression of the tissue, and edges were sealed with additional nail polish. Slides were imaged on a Zeiss LSM 800 confocal microscope using a 40× oil immersion objective. Z-stacks were captured at 512×512 pixel resolution (8-bit), with 2× averaging and 0.27 μm optical sectioning.

#### Volume measurements

Z-stacks were analyzed in Fiji (ImageJ). Volumetric quantification was performed using LabKit (Arzt et al., 2022) and 3D Manager (Ollion et al., 2013) plugins. Neurons were assessed for expression of Calretinin, tdTomato (SERT^Cre^ or Tac1^Cre^), and Tuj1. Type I SGNs were identified via β-tubulin III immunolabeling, which is strongly expressed in type I, but not type II, SGN somata (Lallemend et al., 2007; Barclay et al., 2011; Vyas et al., 2019). As Ai14 mice may exhibit low-level, extranuclear tdTomato expression in the absence of Cre (https://www.jax.org/strain/007914#), we considered nuclear localization of tdTomato as a reliable indicator of Cre-mediated recombination.

### Electrophysiology

#### Preparation

Dissection, dissociation, and culture of SGNs were adapted from a previously described protocol (Iyer et al., 2023). All reagents were obtained from Sigma-Aldrich unless otherwise noted. Temporal bones were isolated in chilled, oxygenated Leibovitz’s L-15 medium supplemented with 10 mM HEPES. The spiral ganglion was dissected from the cochlear epithelium, cleaned of debris and connective tissue, and divided at the midpoint to preserve tonotopic information. Spiral ganglia were digested for 15–20 minutes in L-15 containing 0.05% collagenase, 0.25% trypsin, and 0.05% DNase I at 37°C, then rinsed in L-15 and culture medium. Neurons were dissociated by trituration and plated onto poly-D-lysine-coated coverslips (MatTek). Cells were incubated for 16–24 hours in a bicarbonate-buffered minimal essential medium (MEM; Invitrogen) supplemented with 10 mM HEPES, 1% N2, 2% B27 (Thermo Fisher Scientific), and 1% penicillin-streptomycin. The culture medium was adjusted to pH 7.35 with NaOH and incubated at 37°C in 5% CO₂.

#### Recordings

Recordings were made at 400× magnification using a Zeiss Axiovert 135 TV microscope with Nomarski optics. Signals were recorded and digitized using a MultiClamp 700B amplifier, Digidata 1440A digitizer, and pClamp 10.7 software (RRID:SCR_011323). Pipettes were pulled from borosilicate glass and fire-polished to a resistance of 3–8 MΩ. Pipette capacitance was minimized by wrapping with Parafilm.

Perforated patch recordings used an internal solution containing (in mM): 75 K₂SO₄, 25 KCl, 5 MgCl₂, 5 HEPES, 5 EGTA, and 0.1 CaCl₂, adjusted to pH 7.4 and 270–285 mmol/kg osmolality. Amphotericin B (240 µg/mL) was added fresh from a DMSO stock. Liquid junction potentials (-5.0 mV) were calculated using JPCalc (Barry, 1994) and left uncorrected.

Series resistance (R_s_) was estimated online using MultiClamp software and were not corrected for. Recordings with R_s_ > 35 MΩ were excluded, consistent with prior studies (Ventura and Kalluri, 2019; Bronson and Kalluri, 2023). The average series resistance for SGN recordings was 18.3 ± 7.1 MΩ. Whole-cell capacitance (C_m_) was estimated both online and offline by fitting single-exponential decays of capacitive transients (C_m_ = τ_m_ / R_s_). Input resistance (R_in_) was measured from voltage responses to hyperpolarizing current steps. Recordings were performed at 20-25°C in oxygenated L-15 medium. Only cells forming stable giga-Ohm seals, stable resting potentials, and exhibiting consistent passive properties were included in the analysis.

#### Analysis

All electrophysiology data were analyzed using pClamp 10.7 (Clampfit; MDS Analytical Technologies) and MATLAB R2022b (MathWorks). In current-clamp mode, we measured several intrinsic properties of each neuron, including resting membrane potential, the current threshold for action or graded potentials, and the latency for action or graded potentials. Cells were additionally held near –80 mV using steady hyperpolarizing current injection to standardize resting membrane potential prior to spike initiation and ensure neuronal identity through action potential generation.

In voltage-clamp mode, we measured the approximate steady-state outward current at two time points, 10 ms and 400 ms after stimulus onset, in response to a family of voltage steps. These measurements were taken at both +30 mV (the largest step tested) and –30 mV. Additionally, the maximum inward current within the first 10 ms of the voltage step was recorded.

To estimate current magnitude and voltage dependence, we converted measured currents into whole-cell conductances (*g*_*x*_) by dividing the steady-state current by the driving force for the relevant ion:

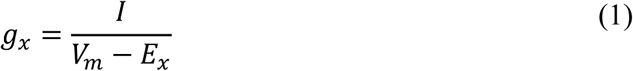

where *I* is current, *V_m_* is the membrane voltage, and *E_x_ i*s the reversal potential for potassium (–81 mV) or sodium (+81 mV), depending on the current type.

Outward conductance at –30 mV was labeled *g*_−30_ and maximal outward conductance at +30 mV was denoted *g*_*max*_. To assess the temporal dynamics of potassium currents, we also calculated *g* ratio defined as the ratio of conductance at 10 ms to that at 400 ms during the +30 mV step.

### Model Framework

Representative type I and type II SGN models were implemented in the Brian 2 simulation environment on Google Colab (code can be here). This model is an adaptation of a Hodgkin-Huxley style framework originally developed in MATLAB to represent vestibular ganglion neurons (Hight and Kalluri, 2016; Ventura and Kalluri, 2019) and was adapted here for spiral ganglion neurons and implemented in the Brian 2 Python Simulator in Google Colab to facilitate flexible implementation and analysis. Briefly, the governing equations were formulated as differential equations in which the net transmembrane current was modeled as the sum of individual ionic and leak currents flowing through parallel conductances (Equation 2).

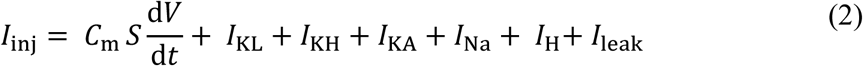

Cell surface area (*S*) times the specific membrane capacitance (*C*m) was set to either be an average of 6 pF; or varied between 5 and 10 pF to represent the range of values across type I and type II SGN. The currents were driven by the following six ionic conductances: sodium (*g_Na_*), low-voltage gated potassium (*g_KL_*), high-voltage gated potassium (*g_KH_*), high-voltage activating and rapidly inactivating potassium conductance (*g_KA_*), hyper-polarization activated mixed-cationic conductance (*g_H_*), and leak (*g_leak_*).

Maximum conductance values (*gKL*, *gKH*, *gKA*, *gNa*) were empirically derived from voltage-clamp recordings of representative neurons: *gKL* was set as the outward conductance at –30 mV (measured at 400 ms), *gKH* was calculated as the difference between the outward conductances measured at +30 mV and –30 mV (both at 400 ms), *gKA* was defined as the difference in conductance at +30 mV between 10 ms and 400 ms.and *g_Na_* was taken as the largest inward conductance within 10 ms of stimulus onset. Reversal potentials were based on expectations for our recording conditions. These values are summarized in **Table 1**.

**Table 1:** Ionic currents used in model simulations with corresponding expressions and comments. Columns indicate the average conductance values in mS/cm^2^ for a standard Type I and Type II SGN, respectively.

| Current | Exp | Type I | Type II | Comments |
| --- | --- | --- | --- | --- |
| $I_{\text{Na}}$ | $\bar{g}_{\text{Na}} m^3 h (V - E_{\text{Na}}), E_{\text{Na}} = -81mV$ | 3.61 | 1.61 | Sodium current |
| $I_{\text{KHT}}$ | $\bar{g}_{\text{KHT}} (n_f n^2 + p(1 - n_f)) (V - E_{\text{K}})$ | 2.09 | 0.5 | High-Threshold Potassium Current |
| $I_{\text{KLT}}$ | $\bar{g}_{\text{KLT}} w^4 z (V - E_{\text{K}}), z = 0.5$ | 2.09 | 0.25 | Low-Threshold Potassium Current |
| $I_{\text{KA}}$ | $\bar{g}_{\text{KA}} a^4 bc (V - E_{\text{K}}), E_{\text{K}} = 81mV$ | 0.065 | 0.69 | In-activating Potassium Current |
| $I_{\text{Ki}}$ | $\bar{g}_{\text{KLT}} w^4 z (V - E_{\text{K}}), z = 1$ | 2.09 | 0.25 | All In-activating KLT Current |
| $I_{\text{HCN}}$ | $\bar{g}_{\text{HCN}} r (V - E_{\text{HCN}}), E_{\text{HCN}} = -41mV$ | 0.91 | 0.91 | Hyperpolarization-activated current |
| $I_{\text{Leak}}$ | $\bar{g}_{\text{Leak}} (V - E_{\text{Leak}}), E_{\text{Leak}} = -65mV$ | 0.03 | 0.03 | Leak current |
| $I_{\text{Cap}}$ | $C * S * dV/dt$ | 6.5 pF | 5.5 pF | Capacitive Current |

The kinetic models governing the channel behaviors are adaptations from a single-compartment vestibular-ganglion model previously described in (Hight and Kalluri, 2016). The equations describing individual ion channel currents were originally based on descriptions from the cochlear nucleus and modified for spiral ganglion neurons.

Each variable’s, *x* ∈ {*m, h, n, p, r, w, z, a, b, c*}, dynamics were modeled using a standard first-order kinetic scheme as follows:

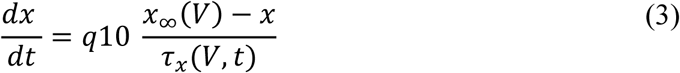

Here:

- *x*_∞_(*V*) is the steady-state activation or inactivation function, defining the voltage-*x*_∞_(*V*)dependent asymptote of the variable.
- *τ*_*x*_(*t*, *V*) *is* the voltage-dependent time constant, determining the rate at which the gating variable approaches *x*_∞_(*V*).
- *q10* accounts for temperature scaling relative to the 22°C reference used in Rothman and Manis.

Both *x*_∞_(*V*)and *τ*_*x*_(*t*, *V*) were defined for each gating variable based on previous models and published data. The specific forms of *x*_∞_ and *τ*_*x*_for each channel are listed in **Table 2** and were taken from, or adapted from, the following sources.

**Table 2:** Steady-state and time constant expressions for gating variables. Voltage V in mV; time constants in ms.

| Channel Type | Steady-State $x_\infty(V)$ | Time Constant $\tau_x(V)$ (ms) |
| --- | --- | --- |
| Na<sup>+</sup> Type I like | $m_\infty = [1 + \exp(-\frac{V+38}{7})]^{-1}$ | $\frac{10}{5e^{\frac{V+60}{18}} + 36e^{-\frac{V+60}{25}}} + 0.04$ |
| | $h_\infty = [1 + \exp(\frac{V+65}{6})]^{-1}$ | $\frac{100}{7e^{\frac{V+60}{11}} + 10e^{-\frac{V+60}{25}}} + 0.6$ |
| K<sup>+</sup> High Threshold | $n_\infty = [1 + \exp(-\frac{V+15}{5})]^{-0.5}$ | $\frac{100}{11e^{\frac{V+60}{24}} + 21e^{-\frac{V+60}{23}}} + 0.7$ |
| | $p_\infty = [1 + \exp(-\frac{V+23}{6})]^{-1}$ | $\frac{100}{4e^{\frac{V+60}{32}} + 16e^{-\frac{V+60}{22}}} + 5$ |
| K<sup>+</sup> Low Threshold | $w_\infty = [1 + \exp(-\frac{V+48}{8})]^{-0.25}$ | $\frac{100}{6e^{\frac{V+60}{6}} + 16e^{-\frac{V+60}{45}}} + 1.5$ |
| | $z_\infty = [1 + \exp(-\frac{V+60}{8})]^{-1}$ | $\frac{1000}{e^{\frac{V+60}{20}} + e^{-\frac{V+60}{8}}} + 50$ |
| K<sup>+</sup> Inactivation | $a_\infty = [1 + \exp(-\frac{V+12}{12})]^{-0.25}$ | $\frac{100}{7e^{\frac{V+60}{14}} + 29e^{-\frac{V+60}{24}}} + 0.1$ |
| | $b_\infty = [1 + \exp(-\frac{V+76}{20})]^{-0.5}$ | $\frac{1000}{14e^{\frac{V+60}{27}} + 29e^{-\frac{V+60}{24}}} + 1$ |
| | $c_\infty = [1 + \exp(-\frac{V+76}{20})]^{-0.5}$ | $\frac{90}{1 + e^{-\frac{66+V}{17}}} + 10$ |
| HCN | $r_\infty = [1 + \exp(\frac{V+95}{7})]^{-1}$ | $\frac{100000}{237e^{\frac{V+60}{12}} + 17e^{-\frac{V+60}{14}}} + 25$ |

### Sodium current (I_Na_)

Sodium conductance was implemented as a classical Hodgkin–Huxley neuronal Na⁺ channel as described in our previous models of vestibular ganglion neurons (Hight and Kalluri, 2016; Ventura and Kalluri, 2019). Activation and inactivation kinetics were based on Rothman & Manis (2003) with modifications to reflect a shift in the activation range as described in previous vestibular ganglion neuron (VGN) models. Sodium conductance density was titrated in simulations to examine the impact of available sodium current on firing patterns. Future improvements in the modeling of sodium current will require measurement methods that better control voltage clamp errors and consideration of persistent and resurgent components (Browne et al., 2017).

### High-threshold K⁺ current (I_KH_) and Low-threshold K current (I_KL_)

Both *I*_KH_ and *I*_KL_ were modeled as using the Rothman & Manis formulation (Rothman and Manis, 2003; Hight and Kalluri, 2016). The steady-state inactivation curve of *I*_KL_ was shifted (via zss=0.5) to match the partially inactivated steady state of *I*_KL_ observed experimentally in SGNs (Mo et al., 2002) and VGNs (Kalluri et al. 2010).

### Transient K⁺ current (I_KA_)

Rather than the slowly activating Kv4-like formulation for the inactivating current described in Rothman & Manis, we implemented a modified transient K⁺ conductance designed to match Kv3.4-like properties as described in dorsal root ganglion (DRG) and SGN (Jagger and Housley, 2003; Ritter et al., 2012; Alexander et al., 2022). Activation was simplified to a three-power gate with a fixed, rapid activation time constant (1.5 ms). Inactivation was modeled with a voltage-dependent time constant and a depolarized steady-state inactivation curve.

### Low-voltage gated completely inactivating current (I_KLi_)

The model parameters are identical to *I*_KLT_ but with the inactivating parameter set to z=1 such that the entire current inactivates. When compared to *I*_KA_, this version of the current tests the relative importance of current inactivation and voltage activation range.

### Hyperpolarization-activated cation current (I_h_)

The model for hyperpolarization-activated cationic current formuation was modified from Rothman and Manis in the following two ways: (1) the half-activation voltage was shifted to more negative potentials (from –76 mV to ∼–96 mV) and the slope factor was slightly adjusted to reflect our measures of HCN channel activation in isolated VGN (Ventura and Kalluri, 2019) and SGN (not shown). Average conductance values were not different between Type I and type II SGN at this age range. Overall small conductance and very negative activation range for HCN conductance meant that the channel did not contribute significant current to the models between resting potential and voltage threshold.

### Leak current (I_leak_)

Leak current followed a simple ohmic form with reversal potential near rest. The leak conductance gleak was held constant across model variants.

In summary, were based on previous models of auditory neurons (Rothman and Manis, 2003b; Jagger and Housley, 2003) as well as dorsal root ganglion neurons for *g_KA_* (Ritter et al., 2015; Alexander et al., 2022). Full equations for conductance density and channel kinetics are provided in Table 1 and 2.

### Statistical analysis

#### Clustering analysis

Dimensionality reduction and clustering analyses were performed using Python code executed in Google Colab. All processing and visualization steps were conducted using standard Python libraries, including pandas, scikit-learn, and umap-learn. 9 basic electrophysical parameters (capacitance, resting membrane potential, spike count, anode break count, latency to peak voltage, current threshold, *g*_*max*_, *g*_*max*_, *and g*_*ratio*_were standardized using z-score normalization with StandardScaler. Uniform Manifold Approximation and Projection (UMAP) was then used to reduce the normalized electrophysiological measurements to two dimensional coordinates (Becht et al., 2019; Pedregosa et al., 2025). UMAP was implemented using umap-learn with parameters set to n_neighbors = 15, min_dist = 0.1, n_components = 2, and random_state = 42 to ensure reproducibility. To determine the appropriate number of clusters, K-means clustering was applied to the UMAP-transformed data across a range of cluster numbers (K = 1–10). The optimal K was estimated using the K that produced the highest silhouette score analysis.

#### Comparison testing

All statistical tests were conducted using OriginPro (OriginLab; RRID:SCR_014212), JMP Pro 17 (SAS Institute; RRID:SCR_014242), Python on Google Colab or Matlab. Sex and tonotopic location were included as factors in initial three-way ANOVAs. If neither factor was significant (*p* > 0.05), data were reanalyzed with one-way ANOVA followed by Tukey’s HSD for post hoc comparisons. Assumptions of equal variance were tested using Levene’s test. If violated (*p* < 0.05), non-parametric Kruskal-Wallis tests were used. Mann–Whitney U tests were employed for pairwise comparisons.

Spearman’s rank correlation was used to test relationships when the dependence between variables appeared nonlinear and the spread in values changed with the dependent variables. Comparisons were made by a permutation test which is amenable to data sets with non-uniform variance across groups (e.g. Fig.7). We used a 2-way ANCOVA to compare the impact of sodium and potassium currents on anode-break spiking (e.g. Fig 9h). The strength of correlations between two continuous variables is reported by Pearson’s r. We used an alpha-level of 0.05 for all statistical tests.

### Code accessibility

The files for the UMAP analysis and the code for the biophysical model will be made available via the ModelDB repository at (tbd).

## Results

### Validation of Cre-reporter lines for biasing recordings toward type II spiral ganglion neurons

Investigating type II spiral ganglion neurons (SGNs) has historically been challenging due to their low abundance within the total SGN (Spoendlin, 1969; Jagger and Housley, 2003; Reid et al., 2004). In the absence of genetic markers or morphological identifiers, only ∼5% of patched neurons are expected to be type II SGNs (Spoendlin, 1969; Nayagam et al., 2011). To improve recording efficiency and confidence in neuron identity, we leveraged genetic strategies to visually identify type II SGNs prior to patch-clamp recordings.

We employed two Cre-driver mouse lines crossed with the Ai14 reporter, which drives tdTomato expression following Cre-mediated recombination. To validate the cell-type specificity of fluorescent labeling, we performed immunohistochemistry for β-tubulin III (Tuj1), a marker selectively expressed in type I SGN somata but absent from type II SGNs (Lallemend et al., 2007; Barclay et al., 2011; Vyas et al., 2019). Additionally, we labeled each sample with antibodies against calretinin (CalB2) which is a calcium binding protein that differentiates type I SGNs by subtype (Petitpré et al., 2018; Shrestha et al., 2018; Sun et al., 2018). Thus, we categorized each cell based on Tuj1 and CalB2 immunoreactivity to infer the identity of each SGN: Tuj1+/CalB2+ (putative pillar-contacting and intermediate type I SGNs), Tuj1+/CalB2– (putative modiolar-contacting type I SGNs), and Tuj1–/CalB2– (putative type II SGNs).

In the first line, SERT^Cre^; Ai14, tdTomato expression was predominantly restricted to type II SGNs. Analysis of whole mount cochlear turns revealed that 85.4% of tdTomato+ cells (117/137 cells from 5 images, 3 mice) lacked Tuj1 immunoreactivity (Figure 1A). Among the 20 tdTomato+/Tuj1+ cells, five also co-expressed Calretinin (CalB2), a marker enriched in medium-and high-spontaneous-rate type I SGNs (Petitpré et al., 2018; Shrestha et al., 2018; Sun et al., 2018). These findings confirm that SERT^Cre^; Ai14 labels the majority of type II SGNs with minimal type I SGN expression, though it misses some type II SGNs located at the cochlear extremes (Vyas et al., 2019).

**Figure 1:**
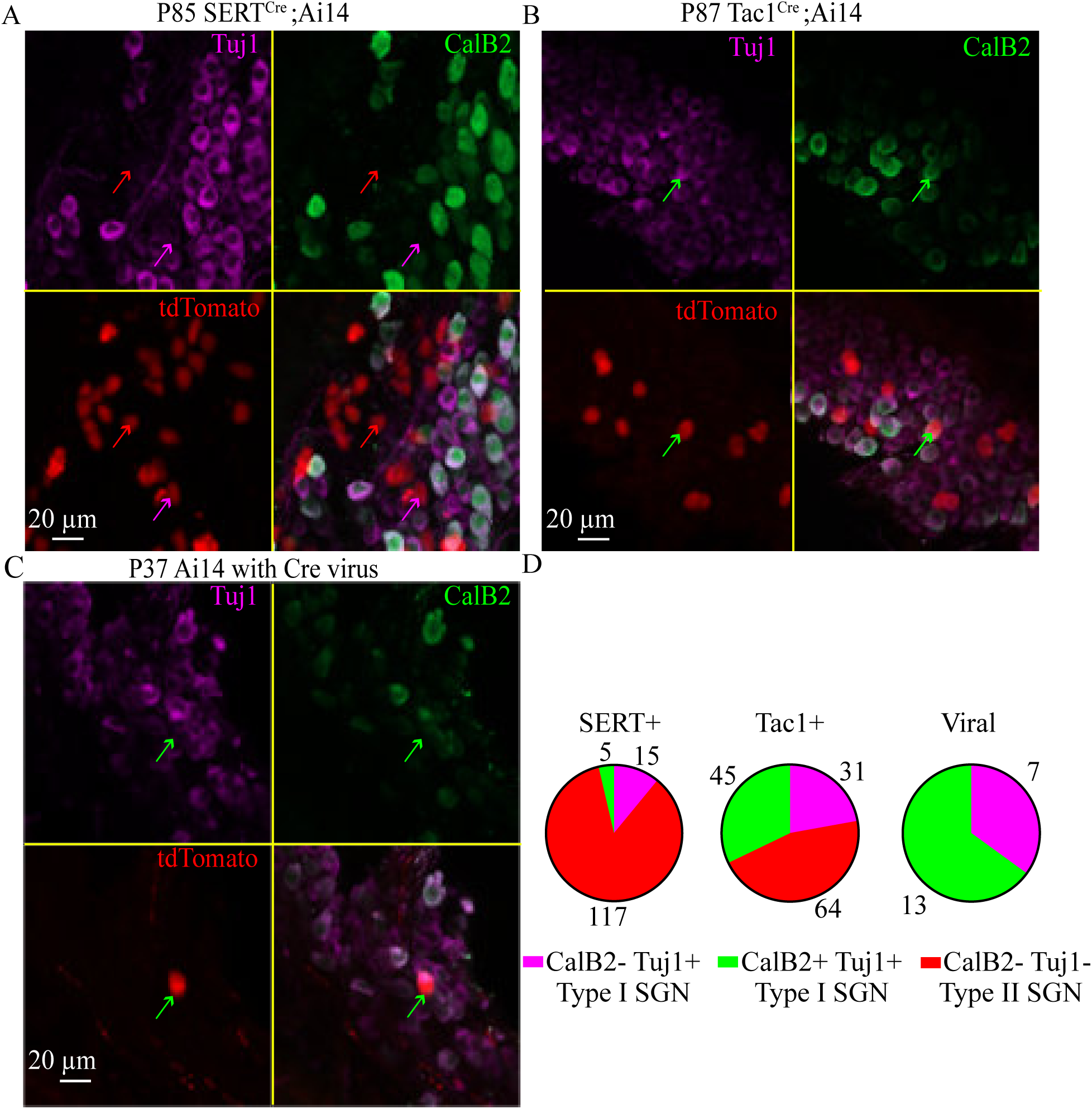
SERTCre;Ai14 and Tac1Cre;Ai14 mice selectively label type II SGNs. 40× confocal images from three genotypes are shown. (A) Image from a P85 SERTCre;Ai14 mouse. (B) Image from a P87 Tac1Cre;Ai14 mouse. (C) Image from a P37 Ai14 mouse with virally delivered Cre. tdTomato is shown in red, Tuj1 (type I SGNs) in magenta, and Calretinin/CalB2 (pillar-contacting type I SGNs) in green. Red arrows mark tdTomato-positive neurons that lack both Tuj1 and CalB2 (putative type II SGNs). Magenta arrows mark tdTomato-positive neurons that co-label only with Tuj1 (type I SGNs, CalB2-negative). Green arrows mark tdTomato-positive neurons that co-labeled with both Tuj1 and CalB2 (CalB2-positive type I SGNs). Scale bar is 20 µm. (D) Pie charts summarize the proportions of SGN subtypes in SERTCre;Ai14 (137 cells, 2 animals), Tac1Cre;Ai14 (140 cells, 4 animals), and virus-injected mice (20 cells, 2 animals).

The second line, Tac1^Cre^; Ai14, induced tdTomato expression in a broader subset of SGNs (Nowak et al., 2021; Sanders and Kelley, 2022). Unlike SERT^Cre^, which favors specificity over inclusivity, Tac1^Cre^ labels a more comprehensive population of type II SGNs but includes a minority of type I SGNs (Figure 1B). In this model, 45.7% of tdTomato+ cells (64/140 cells from 6 images, 4 mice) were also Tuj1+, indicating a substantial fraction of labeled cells were type I SGNs. Among the tdTomato+/Tuj1+ cells, 45 of 76 co-expressed CalB2. CalB2 labels approximately 75% of type I SGNs (Shrestha et al., 2018; Sun et al., 2018; Wang et al., 2023). We verified that the proportion of CalB2 expression in type I SGNs is not affected by tdTomato expression by inducing tdTomato expression stochastically in Ai14 reporter mice through introcerebrovetricular injections of a virus that causes Cre recombinase expression (Figure 1C). Across 3 images from two animals, we observe 13 out of 20 Tuj1+ type I SGNs co-stain for CalB2, similar to previously reported proportions. Overall, we conclude Tac1^Cre^ does not preferentially label specific type I SGN subtypes. In contrast, the relatively low co-expression of CalB2 among SERT^Cre^; Ai14 and Tuj1+ cells suggests a bias toward labeling modiolar-contacting type I SGNs when off-target expression occurs. These distinctions in labeling profiles are summarized in Figure 1D.

Together, these results validate both lines for targeting type II SGNs. SERT^Cre^; Ai14 offers high specificity for type II SGNs while Tac1^Cre^; Ai14 provides an approximately balanced proportion of labeled type I and type II SGNs. These complementary tools enable targeted electrophysiological and morphological characterization of type II SGNs.

### Diversity in the somatic size of type II spiral ganglion neurons

Previous studies have suggested that type II SGNs can be distinguished from type I SGNs by their smaller somatic size (Berglund and Ryugo, 1987; Petitpré et al., 2018). To quantitatively assess somatic size across SGN subtypes, we generated three-dimensional regions of interest (ROIs) for tdTomato-expressing SGNs using the Labkit plugin in FIJI (Figure 2A, Arzt et al., 2022).

**Figure 2:**
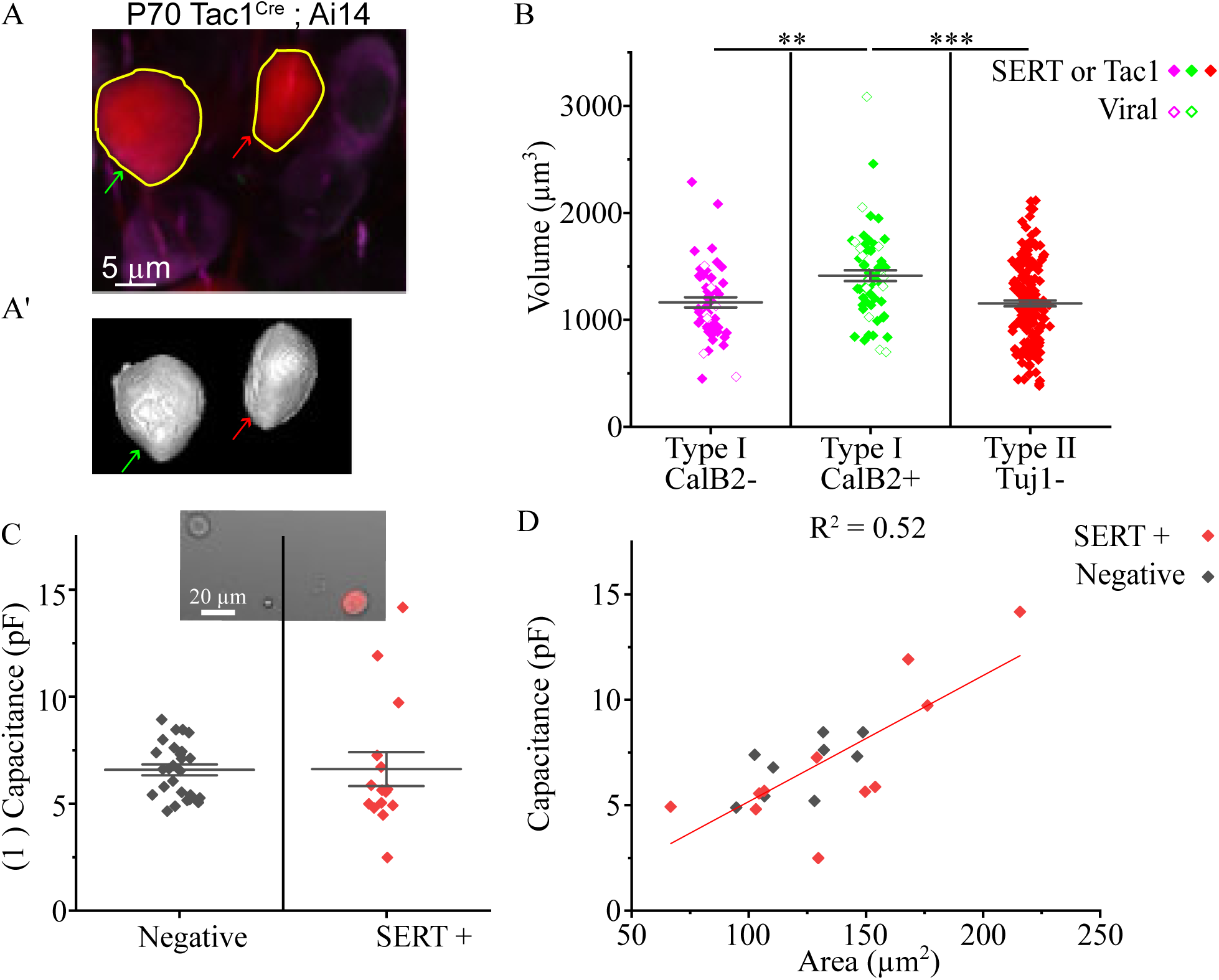
Type II SGNs are not consistently smaller than type I SGNs. (**A**) 40× confocal image of tdTomato+ neurons in a P70 Tac1Cre; Ai14 mouse with hand-drawn ROIs in yellow. Scale bar is 5 µm. (**A’**) Labkit 3D somatic volume reconstructions; red arrow marks putative type II, green arrow marks type I CalB2+. (**B**) Soma volumes of tdTomato+ SGNs: closed symbols are data from Cre lines, open symbols are from viral labeled cells; red marks type II, magenta marks type I SGN not enriched for CalB2 SGN, green marks type I SGN enriched for CalB2+. **0.01 < p < 0.001, ***p < 0.001; error bars = mean ± SEM. (**C**) Capacitance measurements from P5–10 cultured SGNs: black symbols denote cells expressing little tdTomato (putative type I SGNs), red symbols denote cells enriched for tdTomato (putative type II SGNs). Inset: example isolated somata. Scale bar is 20 µm. (**D**) Capacitance vs. cross-sectional area with linear fit.

Somatic volume significantly differed across tdTomato+ cells that lacked Tuj1 and CalB2, labeled for Tuj1 but not CalB2, and labeled for both Tuj1 and CalB2 (Kruskal-Wallis ANOVA, H(2) = 21.55, *p* = 2.09 × 10⁻⁵). Tuj1+/CalB2+ cells had the largest volumes (1417.09 ± 50.39 μm³), which were significantly greater than both Tuj1+/CalB2– (1167.33 ± 47.56 μm³) and Tuj1–/CalB2–SGNs (1157.91 ± 27.26 μm³) (Dunn’s post hoc test, *p* = 0.0013 and *p* = 1.93 × 10⁻⁵, respectively; Figure 2B). However, Tuj1+/CalB2– and Tuj1–/CalB2– SGNs did not differ significantly in size (*p* = 1.0). These results indicate that only CalB2+ type I SGNs exhibit significantly larger somata, while type II SGNs closely match the size of CalB2– type I SGNs.

To further validate this observation, we measured somatic volumes in SGNs from two Ai14 reporter mice injected with a virus encoding Cre recombinase. This strategy induces sparse, stochastic labeling across the SGN population, independent of SERT^Cre^ or Tac1^Cre^ expression. The somatic volumes of these virally labeled, Tuj1+ SGNs (n = 20 cells; 1343.97 ± 129.33 μm³) were not significantly different from those of Tuj1+ SGNs labeled in SERT^Cre^ or Tac1^Cre^ mice (n = 94; 1294.43 ± 35.59 μm³; two-way ANOVA, *F*(1,112) = 0.042, *p* = 0.84, Tukey’s post hoc test). Thus, labeling through genetic or viral methods did not affect somatic size estimates, reinforcing the conclusion that CalB2– type I SGNs and type II SGNs are comparable in size.

We also assessed SGN size following dissociation and plating on poly-D-lysine-coated coverslips for electrophysiological recordings. Differential interference contrast (DIC) imaging revealed that SERT^Cre^ + (putative type II) SGNs had cross-sectional areas ranging from 64.9 to 215.81 μm² (mean = 134.64 ± 7.57 μm², *n* = 27, data not shown), while SERT^Cre^ – (putative type I) SGNs ranged from 55.0 to 250.24 μm² (mean = 125.81 ± 4.33 μm², *n* = 60, data not shown). Although SERT^Cre^ + cells showed greater variance and included both the smallest and largest recorded cells, the average difference between groups did not reach statistical significance (Mann-Whitney U = 686, Z = –1.13, *p* = 0.26).

Among the recorded neurons, SERT^Cre^ + SGNs exhibited a broader range of cell sizes than their SERT^Cre^ – counterparts, which clustered closer to the mean. For these same neurons, we measured membrane capacitance as an indirect indicator of cell size. Capacitance values were similar between SERT^Cre^ + (6.61 ± 0.79 pF) and SERT^Cre^ – (6.58 ± 0.25 pF) SGNs (Mann-Whitney U = 237, Z = 1.12, *p* = 0.26; Figure 2C). Furthermore, capacitance correlated linearly with cross-sectional area (linear fit, *R²* = 0.52, *df* = 23, *p* = 3.14 × 10^-5^; Figure 2D), validating the use of capacitance as a proxy for cell size in this context.

Together, these data reveal that while SERT^Cre^ + type II SGNs span a broader range of somatic sizes, their average size is comparable to that of CalB2– type I SGNs. Thus, soma size alone is insufficient to reliably distinguish type I and type II SGNs, particularly when CalB2 expression is absent.

### Diverse Intrinsic Firing Properties Among Spiral Ganglion Neuron Subtypes

To investigate the intrinsic excitability of SGN subtypes, we performed perforated patch-clamp recordings from dissociated and cultured neurons harvested from SERT^Cre^; Ai14, Tac1^Cre^; Ai14, and control mice (P5–P10). To facilitate access to the neuronal membrane, SGNs were cultured overnight to promote detachment of the overlying satellite cells. In the absence of afferent synaptic connections in vitro, we identified putative type II SGNs based on tdTomato fluorescence in SERT^Cre^ + and Tac1^Cre^ + mice, while non-fluorescent neurons from reporter mice and neurons from wild-type mice served as putative type I SGN controls. In total, we recorded from 15 SERT^Cre^ + SGNs, 9 SERT^Cre^+ SGNs, 26 non-fluorescent SGNs from reporter mice (“negative”), and 12 SGNs from wild-type mice (“control”).

In current-clamp mode, we injected depolarizing current steps ranging from –60 pA to +330 pA and assessed various features of the resulting voltage responses. A primary classification was whether neurons produced action potentials. Most neurons in the negative group (23/26), representing putative type I SGNs, fired a single action potential at the onset of depolarizing current injection (Figure 3A,C). Action potentials were identified based on the presence of a sharp voltage upstroke, indicating rapid activation of voltage-gated sodium channels. The remaining three neurons in this group exhibited graded depolarizations that scaled with current amplitude, lacking the rapid rise typical of sodium-driven spikes.

**Figure 3:**
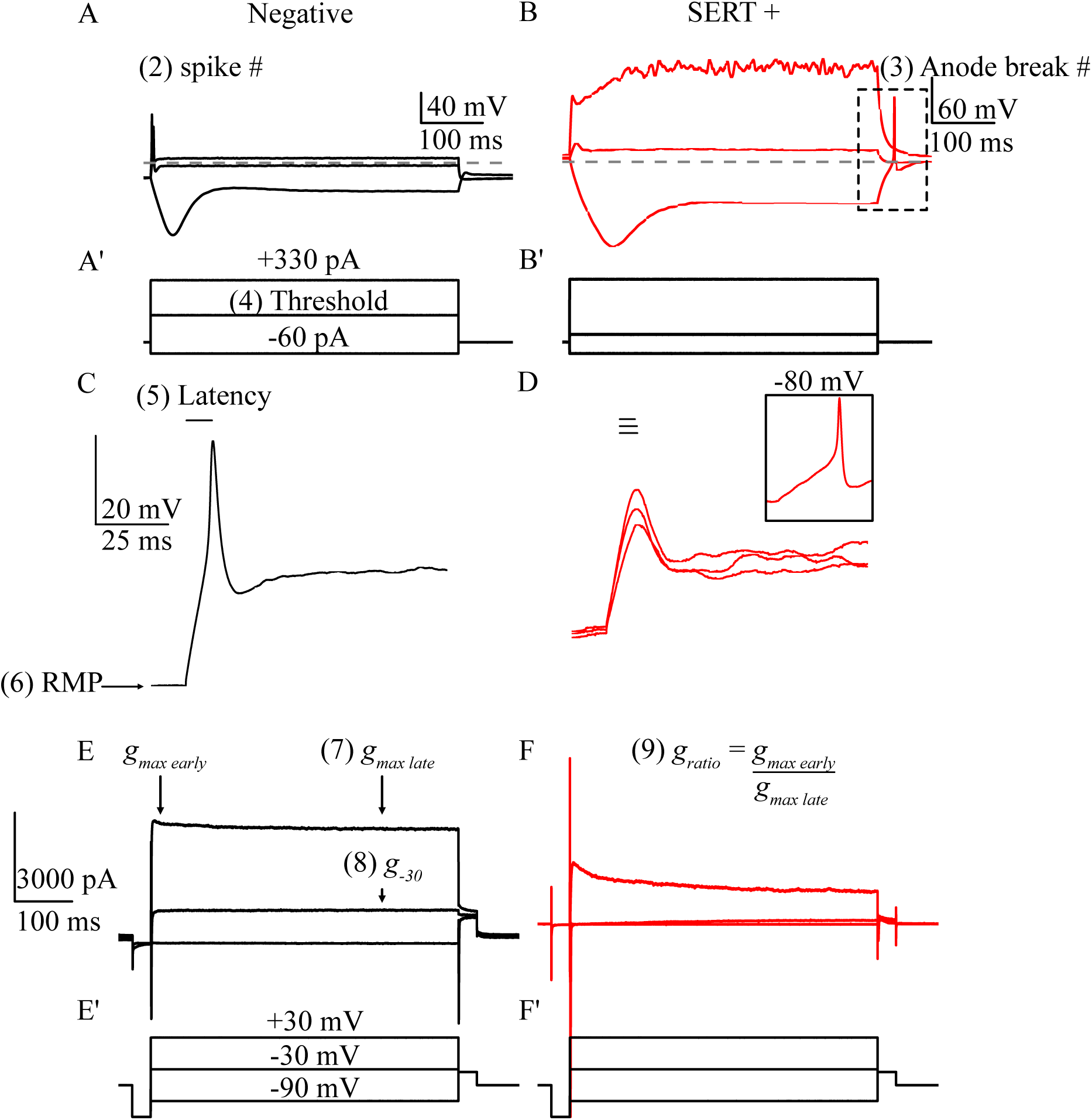
Type I and type II SGNs differ in current and voltage responses. (A–B) Current-clamp traces: tdTomato-(putative type I SGNs, A) and SERT+ (putative type II SGNs, B). (A’–B’) Stimulus protocols. (C-D) Threshold response zoom for same negative neuron as A (C) and threshold response and next two suprathreshold responses for the same SERT+ neuron as B (D). latency = time to peak; inset shows action potential when cell is held at -80 mV. (E–F) Voltage-clamp responses for same negative neuron as A (E) and SERT+ neuron as B (F); *g* = outward conductance at specified voltages and times. (E’-F’) Voltage step protocols.

By contrast, the majority of SERT^Cre^ + SGNs (11/15) failed to generate action potentials from rest, even at the highest current injections tested (Figure 3B,D). These neurons instead displayed graded voltage responses (Figure 3D). However, hyperpolarizing the membrane to –80 mV relieved apparent sodium channel inactivation and permitted spike generation (Figure 3D, inset), consistent with voltage-dependent inactivation at resting membrane potential (Markowitz and Kalluri, 2020). Additionally, four of these SERT+ neurons produced rebound spikes, so-called anode break spikes, upon termination of hyperpolarizing current steps. These anode break spikes were observed exclusively in SERT^Cre^ + neurons and one Tac1^Cre^ + neuron; no anode break spikes were seen in negative or control neurons. A minority of SERT^Cre^ + neurons (4/15) did produce action potentials from rest, and two of these also exhibited anode break spikes (Figure 2D,H). Additionally, one SERT^Cre^ ^+^ SGN and one Tac1^Cre^ + SGN were the only two cells to fire two or more action potentials to depolarizing current at rest.

Together, these results suggest that while type I SGNs typically exhibit a consistent firing pattern, a single spikes in response to depolarization, type II SGNs display greater diversity, including non-spiking, anode-break spiking, and multiple firing phenotypes. However, spiking behavior alone was insufficient to clearly distinguish SGN types. Graded responses were observed in a minority of type I SGNs, and anode-break spiking, though specific to putative type II SGNs, was relatively rare.

To improve subtype classification, we next quantified additional intrinsic properties. In current clamp, we measured the current threshold for spike or graded response generation, the latency from stimulus onset to peak voltage at threshold, and resting membrane potential (RMP). To further characterize membrane properties, we recorded voltage-clamp responses to a family of voltage steps from –90 mV to +30 mV in 5 mV increments, allowing us to estimate potassium conductance (Figure 3E-F). Steady-state conductance (*g*) was measured 400 ms after the voltage step for –30 mV (*g_-30_*) and +30 mV (*g_max_*) to reflect low- and high-voltage-activated potassium channels, respectively. Additionally, the ratio of conductance at 10 ms and 400 ms after the onset of the +30 mV step (*g* ratio) was used to assess temporal dynamics of outward current.

Ultimately, no single biophysical parameter reliably distinguished type I from type II SGNs. Substantial variability within each group meant that none of these features alone could serve as a definitive marker for SGN subtype identity.

### Biophysical clustering reveals distinct SGN subtypes

As shown in Figure 1D, genotype-based classification using SERT^Cre^ or Tac1^Cre^; Ai14 is imperfect, as neither Cre line exclusively labels type II SGNs. This raises the possibility that the observed variability in intrinsic properties among SERT^Cre^+ and negative (tdTomato–) neurons may reflect a mixture of both type I and type II SGNs within each group.

To address these limitations, we used an unsupervised clustering approach based on a comprehensive set of biophysical properties measured in current- and voltage-clamp modes. We compiled nine normalized parameters and applied Uniform Manifold Approximation and Projection (UMAP) to visualize similarity across neurons in two dimensions. These parameters included membrane capacitance (Figure 2C), spike count during depolarization, presence of anode break spikes, current threshold, latency to peak voltage, resting membrane potential, and three voltage-clamp conductance measures: *g_-30_*, *g_max_*, and the *g* ratio (Figure 3). Genotype and tdTomato expression were excluded from the clustering analysis.

UMAP analysis of 62 high-quality recordings revealed two primary clusters, identified via K-means clustering (Figure 4A). These clusters corresponded well with tdTomato expression: SERT^Cre^ + neurons tended to group into cluster 2, while negative and control SGNs mostly grouped into cluster 1 (Figure 4B). Spiking phenotype was a strong predictor of cluster identity (Figure 4C). Neurons in cluster 2 typically did not fire action potentials in response to depolarizing current. Notably, this cluster also separated by the presence of anode break spikes, suggesting a possible subcluster of anode-break spiking neurons.

**Figure 4:**
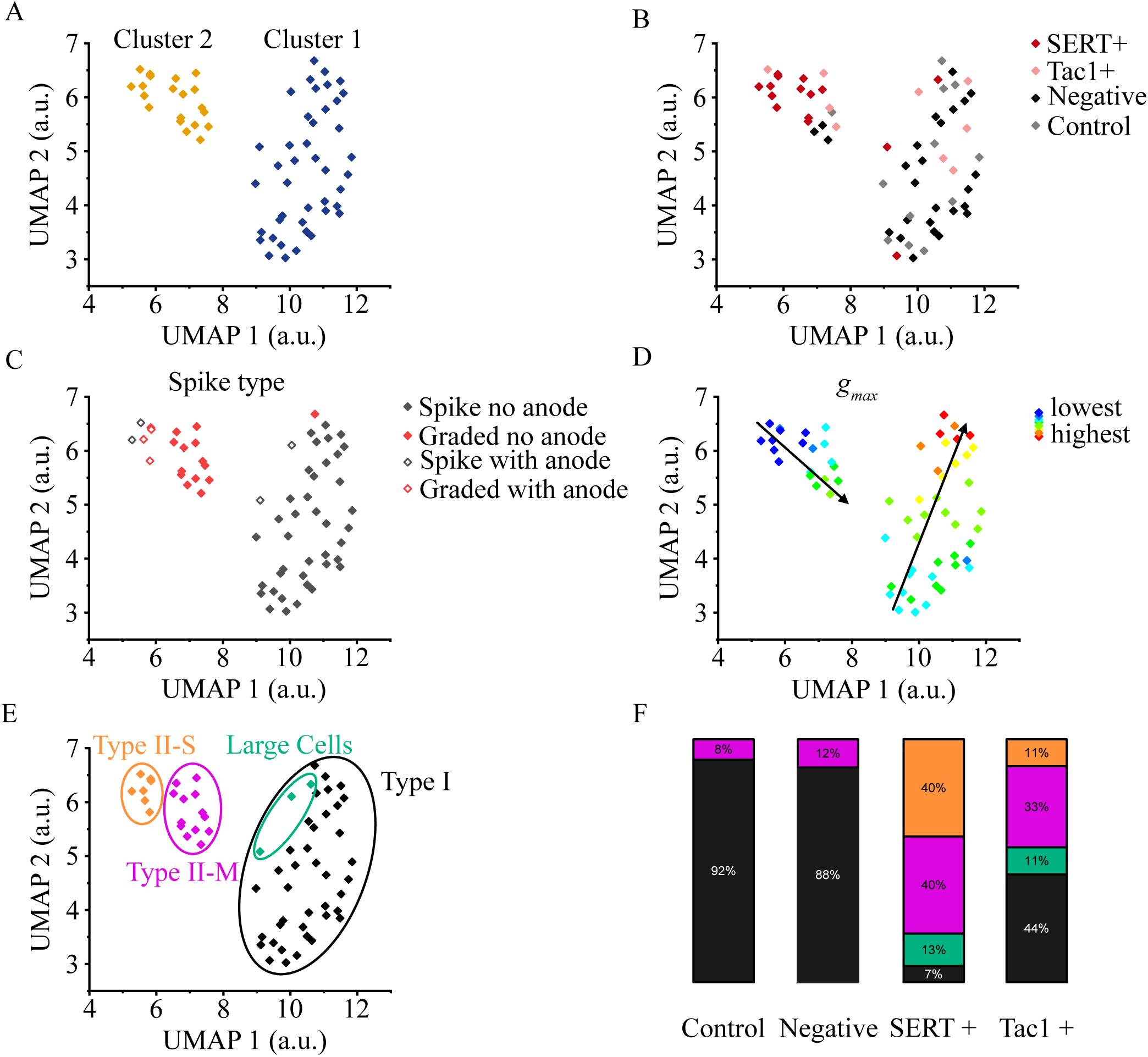
UMAP clustering segregates SGNs by subtype. (A–E) Scatterplots of UMAP scores. (A) Scatterplot colored by cluster. Blue points indicate individual cells in cluster 1 (putative type I SGNs), and gold points indicate cells in cluster 2 (putative type II SGNs). (B) Points colored by genotype/tdTomato expression. Dark red denotes SERT+, pink denotes Tac1+, black denotes tdTomato−, and gray denotes cells randomly drawn from non-fluorescent control animals. (C) Points colored according to spiking behavior. Gray denotes cells that fire an action potential in response to a current injection from rest, red denotes cells that do not fire from rest, and unfilled symbols indicate cells that fire anode break spikes. (D) Scatterplot colored according to the value of g_max. Arrows highlight the gradient of g_max from lowest to highest. (E) Scatterplot identifying subtypes within the type II cluster. Orange denotes type II-S SGNs, magenta denotes type II-M SGNs, green denotes large cells, and black denotes type I SGNs. (F) Bar plot showing subtype distribution by genotype.

Within cluster 1, *g_max_* values showed a continuous gradient along the cluster’s vertical axis (Figure 4D). This distribution likely reflects known biophysical differences among type I SGN subtypes, where higher *g_max_* values are associated with modiolar-contacting and lower *g_max_* with pillar-contacting SGNs (Markowitz and Kalluri, 2020). Although we did not assess synaptic position directly, the alignment suggests the clustering approach captured meaningful subtype variation. A similar gradient was observed in cluster 2, suggesting potential subtype heterogeneity among type II SGNs as well. As gradients in biophysical properties have been observed in SGNs depending on cochlear position (Adamson et al., 2002) we also looked for differences between cells originating from apical or basal portions of the cochlea. Either by looking at individual parameters or on UMAP plots we found no clear differences between apical or basal cells in our dataset (data not shown). Likewise, repeating this analysis for cells from male or female mice also had no obvious organization (data not shown).

Taken together, these results support the interpretation that cluster 1 corresponds to type I SGNs. Similarly, cluster 2, enriched for SERT^Cre^ + and Tac1^Cre^ + neurons, likely corresponds to type II SGNs. Substructure within cluster 2 suggests multiple type II subtypes. Based on capacitance and spiking behavior, we tentatively define two subgroups: type II-S SGNs, smaller cells exhibiting anode break spikes, and type II-M SGNs, more moderately sized cells with graded responses but no anode break spikes (orange and magenta, respectively, Figure 3E).

A third group of three large tdTomato+ neurons (green, Figure 3E) occupied an intermediate position between clusters 1 and 2. These neurons had the largest measured capacitance and soma area (Figure 1F–G), and 2 of 3 displayed both evoked and anode-break spiking. Given their unique properties and low sample size we will simply refer to them as large cells and will exclude them from statistical comparisons.

The distribution of neurons across clusters closely matched expectations from immunohistochemistry (Figure 3F). Specifically, 11 of 12 control SGNs (92%) and 23 of 26 negative SGNs (88%) were assigned to cluster 1, consistent with the ∼90-95% prevalence of type I SGNs in the cochlea. Conversely, only 1 of 13 non-large SERT^Cre^ + neurons (8%) was assigned to cluster 1, aligning with prior estimates that ∼15% of SERT^Cre^ + neurons co-label with the type I marker Tuj1 (Figure 1C). Tac1^Cre^+ neurons, which showed ∼55% Tuj1 co-labeling, were approximately evenly split with 4 of 8 non-large cells (50%) falling into cluster 2.

Figure 5 shows the individual breakdown for all of the 62 cells based on their relative rankings for each parameter than informed the UMAP analysis. Cells are ranked by their UMAP1 score along the x-axis such that putative type II SGNs appear on the left-hand side and putative type I SGNs appear on the right-hand side (Figure 5A). Secondly, each block is colored based on the whether the cell was closer to the minimum (bluer colors) or to the maximum (redder colors) for that parameter. For example, the color code illustrates that certain parameters, such as threshold, smoothly change across the UMAP1 axis suggesting a correlation between some electrophysiological features and SGN type identity. Other parameters such as spike number, which is more binary in nature, have more sudden transitions. Although no single feature alone perfectly delineates type I from type II SGNs, analyzing features in conjunction reveals patterns that align well with SGN identity.

**Figure 5:**
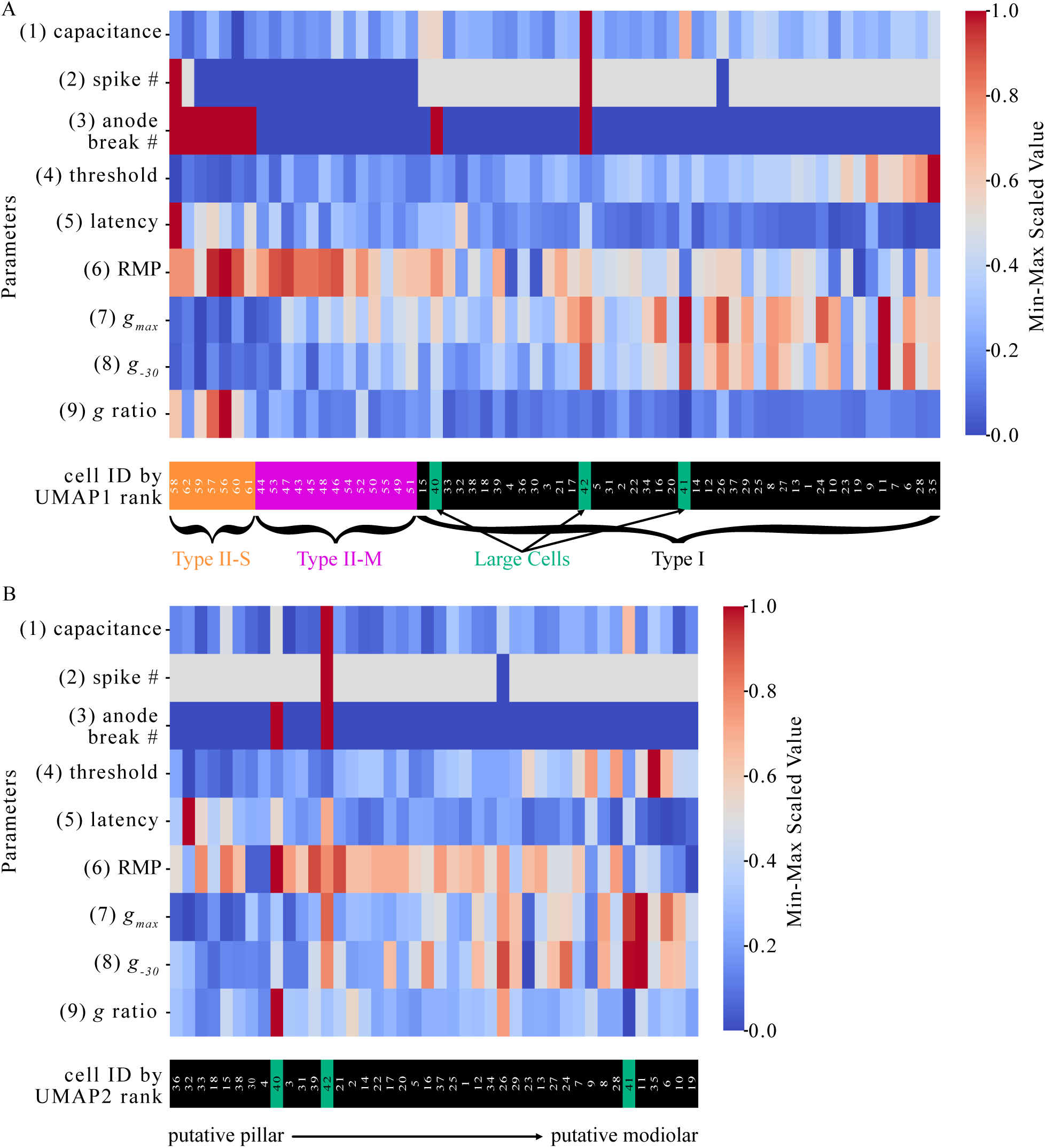
Heatmaps of biophysical parameters contributing to UMAP clustering. (A) Heatmap of all cells ordered by UMAP1 rank (left = lowest, right = highest). (B) Heatmap of cluster 1 ordered by UMAP2 rank (left = lowest, right = highest). Orange marks type II-S SGN, magenta marks II-M SGN, green marks large cells, black marks type I SGN. Cool colors equal low values, warm colors equal high values.

Furthermore, within cluster 1 (type I cluster) parameters such as current threshold, latency, and *g_max_* gradually vary along the UMAP2 axis (Figure 5B). These gradients in biophysical properties align with previously described variations within type I SGNs based on the putative modiolar to pillar location of their synapses with inner hair cells (Markowitz and Kalluri, 2020). This suggests that the UMAP scores for each cell capture relevant factors for characterizing all sub-groups of SGNs. Specifically, UMAP1 appears to divide SGNs into either type I or type II groups while UMAP 2 identifies variation within each SGN type.

In summary, unsupervised clustering of electrophysiological data effectively distinguished type I and type II SGNs, recapitulated biophysical diversity within type I SGNs, and revealed previously unappreciated diversity within the type II population. This approach provides a functional framework for SGN classification that complements and extends genetic labeling strategies.

### Outward currents are dominated by an inactivating component in type II but not type I SGNs

One of the most striking features of the putative type II-S SGN subtype was the relatively large *g* ratio. To further characterize the source of this inactivating potassium currents, we adapted a voltage protocol previously used by Jagger and Housley (2003). Cells were held first at –100 mV (Figure 6A, black trace) for 0.5 seconds, followed by a +40 mV depolarization for 1 second. The protocol was then repeated with the prepulse instead at –40 mV (Figure 6A, cyan trace). The –40 mV prepulse inactivates transient conductances, while the –100 mV prepulse relieves inactivation. The resulting difference (Figure 6B, gray trace) isolates the inactivating component of the outward current.

**Figure 6:**
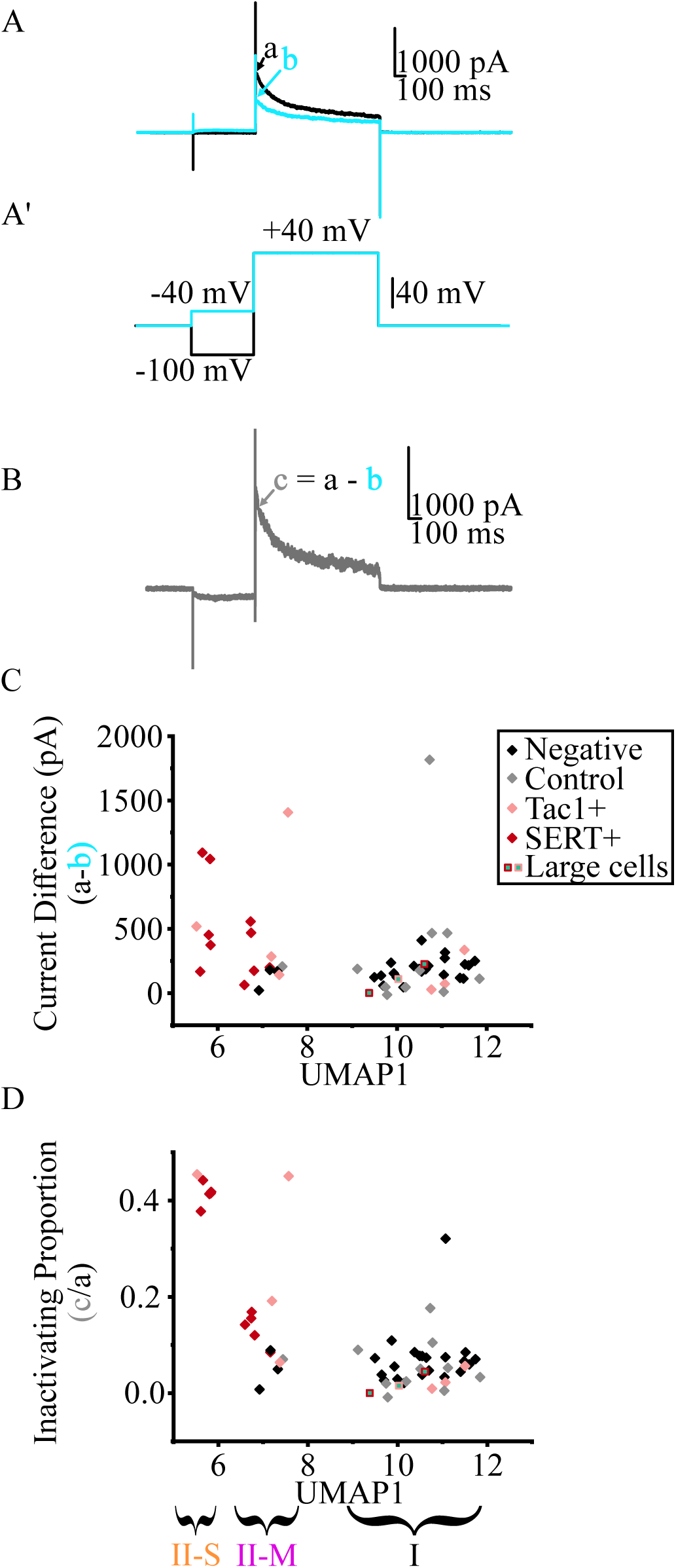
Type II SGNs express an inactivating outward current. (A) Voltage-clamp example for a type II-S SGN; black line corresponds to a response following a −100 mV preconditioning step, cyan line corresponds to a response with a −40 mV preconditioning step. (A’) Voltage step protocol. (B) Difference current (black minus cyan). (C–D) Scatterplots of current differences (C) and inactivating proportion of total current (D) vs. UMAP1 score. Colors: black = tdTomato−, gray = control, pink = Tac1+, red = SERT+. Squares with green centers = large cells.

Although type II-S SGNs had the smallest total outward currents, they exhibited greater inactivating currents than type I SGNs. Type II-M had similar total amounts of inactivating current whereas type I SGNs mostly had minor inactivating currents despite their relatively larger total outward current values (Figure 6C). In type II-S SGNs (n = 6), the inactivating current comprised ∼42% of the total outward current following the –100 mV prepulse. This proportion was much higher than in type I SGNs (n = 34), where inactivation was minimal. Type II-M SGNs (n = 13) also appear to have a more moderate proportion of inactivation in between type II-S SGNs and type I SGNs (Figure 6D). These results mirror earlier observations where *g* ratio values suggested pronounced inactivation mostly in type II-S SGNs, some in type II-M SGNs, and very little in type I SGNs.

Interestingly, not all type II SGNs displayed strong inactivating currents with several type II-M cells exhibiting minimal inactivation. Nonetheless, the presence of a robust inactivating potassium current remains a distinguishing and consistent feature of the type II-S subtype.

### Type II SGNs also have smaller sodium currents than type I SGNs

Given the marked differences in excitability and potassium conductances among SGN subtypes, we next examined whether sodium conductances also differ between type I and type II SGNs. Using voltage-clamp recordings, we quantified peak inward sodium currents immediately following the onset of depolarizing voltage steps (Figure 7A). These inward currents rapidly reached a maximum amplitude and then quickly inactivated. As expected, sodium current amplitude increased with depolarization up to a peak, followed by a decline as the membrane potential approached the sodium reversal potential.

**Figure 7:**
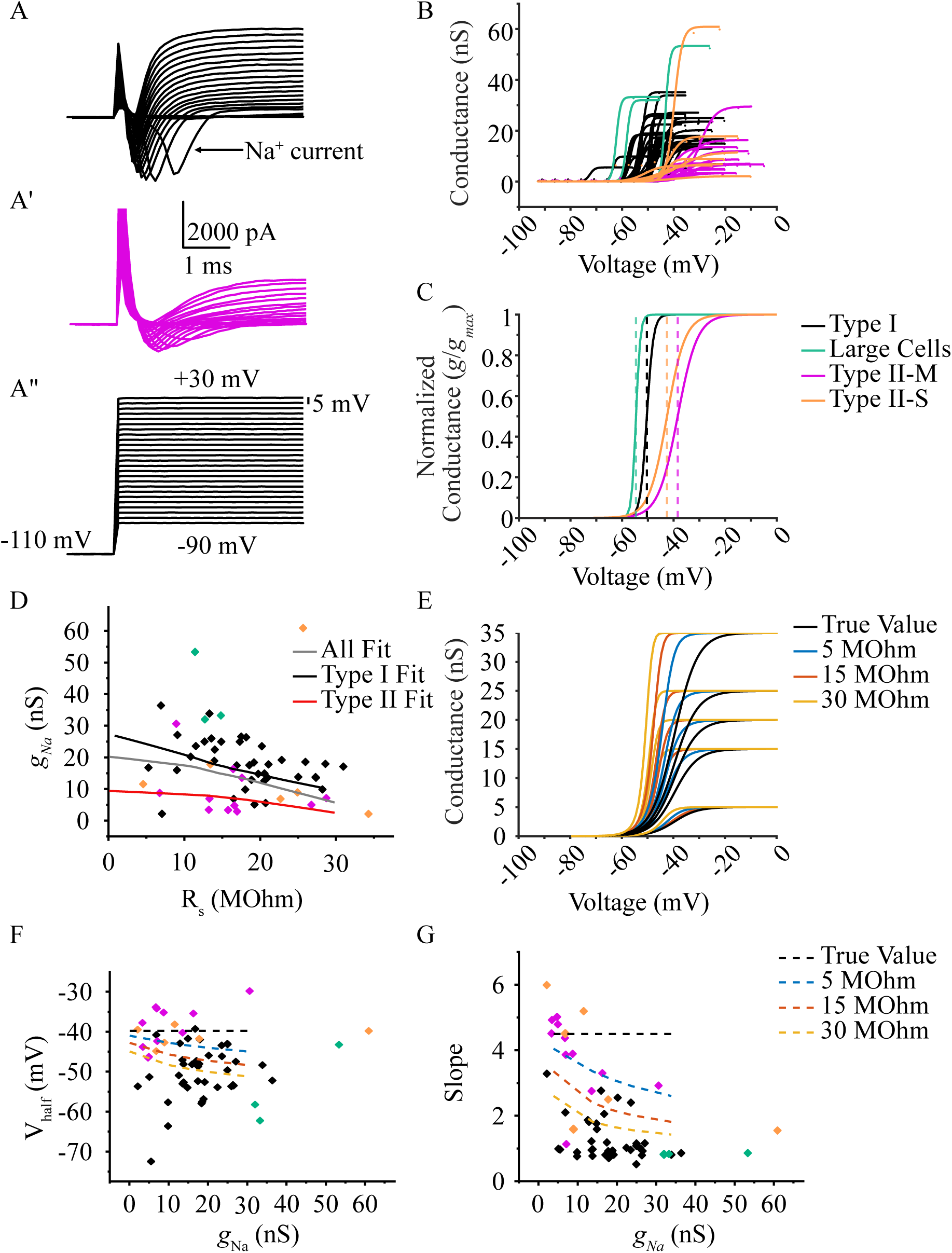
Type II SGNs have reduced sodium conductance. ((A–A′) Example voltage-clamp traces from a type I SGN (A) and a type II-M SGN (A′). (A″) Voltage-step protocol. (B–C) Boltzmann fits of *g_Na_* with normalization (B) and without normalization (C). Colors: black, type I SGNs; green, large cells; magenta, type II-M SGNs; orange, type II-S SGNs. (D) *g_Na_* plotted against series resistance (R_s_). Fitted lines for all SGNs (gray), type I SGNs (black), and type II SGNs (red). (E) Model illustrating the effect of R_s_ on *g_Na_* Boltzmann activation curves. Black, blue, orange, and yellow lines correspond to R_s_ values of 0, 5, 15, and 30 MΩ, respectively. (F–G) V_half and slope plotted as a function of *g_Na_*. Dashed lines show the model-estimated effects of R_s_.

To characterize the voltage dependence of sodium current activation, we fitted Boltzmann functions to the sodium conductance across voltage steps for each cell (Figure 7B). From these fits, we extracted the voltage at half-maximal activation (V½), the slope factor, and the maximum sodium conductance. We then averaged these parameters across cells within each SGN subtype to generate representative activation curves (Figure 7C).

Type I SGNs exhibited a wide range of *g_Na_* values but, on average, had larger sodium conductance than type II-S or type II-M SGNs. Aside from a single outlier, type II-S neurons had *g_Na_* values similar to Type II-M neurons, so we pooled these subtypes for subsequent analyses. Because series resistance (R_s_) can obscure measurements of sodium conductance, we asked whether the apparent subtype differences were better explained by R_s_ or by SGN identity. We observed a weak but statistically significant monotonic relationship between *g_Na_* and R_s_ (Spearman ρ = –0.259, p < 0.05; Figure 7D).

To determine whether type II SGNs truly have smaller *g_Na_* than type I SGNs after accounting for R_s_, we performed a non-parametric permutation test. Group labels (type I vs. type II) were randomly shuffled 10,000 times, and the mean difference in *g_Na_* was recalculated for each shuffle to generate a null distribution. The observed difference between subtypes (–7.77) was significant relative to this null distribution (p = 0.0022). Thus, the reduction in *g_Na_* in type II SGNs cannot be explained by voltage-clamp artifact and likely reflects a genuine subtype difference in sodium conductance.

We also observed an overt rightward shift in the V_½_ of sodium activation and shallower slopes in type II SGNs relative to type I SGNs, however these effects may also be influenced by R_s_. To account for the effects of R_s_, we simulated neurons with varying *g_Na_* but identical true V_½_ and slope values. We then fit Boltzmann curves to the *g_Na_* data after introducing the distortions expected from series resistance (Figure 7E). This approach allowed us to examine how series-resistance artifacts could create apparent correlations between *g_Na_*, V_½_, and slope. Our simulations show that series resistance can make the measured V_½_ appear more hyperpolarized, with a larger effect in cells with higher *g_Na_*, such as the case of most type I SGNs (Figure 7F). Similarly, the slope of the activation curve appears steeper than it truly is with increasing R_s_ (Figure 7G). Therefore, while the differences in the size of *g_Na_* are probably real, the covariation of V_½_ and slope with *g_Na_* may be partly influenced by a voltage-clamp artifact. Thus, while we cannot be confident that sodium channel compositions differ between type I and type II SGNs, but the results suggest that type II SGNs have lower sodium current density.

### Type II SGNs have a reduced capacity to respond to repetitive stimulation

Potassium channels are critical for regulating repetitive neuronal firing by shaping afterhyperpolarizations and promoting sodium channel recovery from inactivation (Iwasaki et al., 2008; Kalluri et al., 2010; Cao and Oertel, 2011; Hight and Kalluri, 2016). Given the observed differences in potassium current magnitude and kinetics between SGN subtypes, we hypothesized that type II SGNs would be less capable of following high-frequency stimulation, especially given their reduced sodium conductance.

To test this, we delivered trains of 3 ms long square current pulses at frequencies of 6, 10, 20, and 100 Hz for 0.5 seconds, increasing stimulus amplitude up to 450 pA in 25 pA steps. At stimulation frequencies less or equal than 20 Hz, any SGN capable of firing an action potential at rest reliably spiked to all pulses (data not shown). However, at 100 Hz, only type I SGNs could maintain consistent firing. Specifically, 21 out of 28 type I SGNs fired an action potential in response to every one of the 50 pulses. In contrast, all but one type II SGN failed to produce more than a single action potential per train, instead generating graded responses

Next, we compared the response to high frequency stimulation when all cells were capable of firing at least one action potential by holding the cells at -80 mV (Figure 8A-B). The differential ability of type I and type II SGNs to follow high-frequency stimulation persisted when all of the cells were held at -80 mV to relive the sodium channel inactivation of type II SGNs. In this case, 9 out of the 11 putative type II SGNs tested fired 1 total spike and the remaining two fired 2 total spikes. Type I SGNs maintained a similar pattern as compared to baseline with 11 out 15 cells firing to all 50 current impulses (Figure 8D).

**Figure 8:**
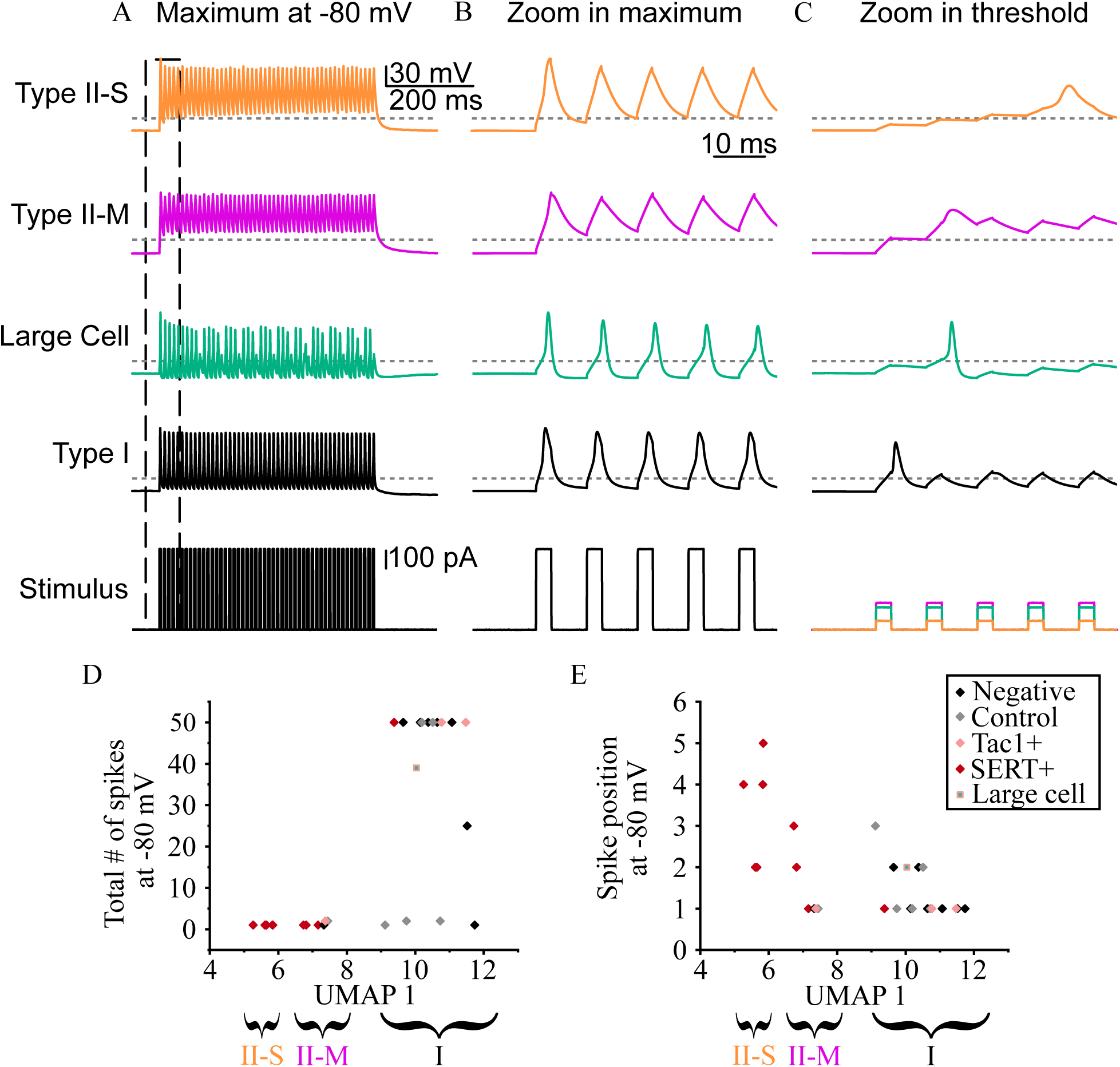
Type I SGNs sustain high-frequency firing but type II SGNs do not. (A–D) Voltage response examples to suprathreshold100 Hz, 3 ms current steps when cells are held at -80 mV; in order from top to bottom type II-S SGN, type II-M SGN, large cell, type I SGN, and stimulus protocol. B. Zoom of first five responses to suprathreshold currents in A. C. First five spikes at action potential threshold when held at −80 mV. (D-E) Total spike counts at 100 Hz (F) and position of the pulse to first initiate spike at -80 mV (G) vs. UMAP1 rank. Colors: black symbols denote tdTomato negative cells, gray denotes control cells, pink denotes Tac1 positive cells, red denotes SERT positive cells. Squares with green center denote large cells.

Although most type I SGNs fired on the first current impulse, many type II SGNs showed delayed spiking even when held at -80 mV (Figure 8C). Of the 11 type II SGNs tested at -80 mV, 7 failed to spike until the second or later stimulus, and all 4 that fired immediately were classified as type II-M SGNs. Strikingly, the majority of tested type II SGNs fired on or after the fourth current impulse (Figure 8E). In these 3 type II-S SGNs the current impulses build in a step-like pattern where there is little loss of signal until the spike occurs (Figure 8C top). Altogether, these findings reinforce earlier observations from extended square-pulse stimulation (Figure 3) that type II SGNs exhibit reduced excitability and integrate current to reach firing threshold, a behavior incompatible with high-frequency spiking. In contrast, type I SGNs are well-suited to high-frequency firing, likely due to more robust and faster potassium currents that repolarize the membrane and support sodium channel recovery between pulses.

### Simulated SGNs reproduce key features of real SGN electrophysiological recordings

Although SGNs display diverse responses to current injection, the specific features that define the characteristic firing behavior of each SGN type remain unclear. As shown in Figure 6, an inactivating outward potassium conductance (*g_KA_*) is prominently expressed in most type II SGNs but are largely absent in type I SGNs. Type II-M and type II-S SGNs also exhibit reduced outward conductance density, both near the action potential threshold (*g_KL_*) and at more depolarized voltages (*g_KH_*), compared to type I SGNs.

Disentangling the contributions of these individual conductances is difficult using pharmacological methods due to the limited specificity of available potassium channel blockers (Wulff et al., 2009). To address this, we adapted a conductance-based model of inner ear neurons (Hight and Kalluri, 2016) to simulate how neurons will respond to electrical inputs. The model functions by calculating and summing the current that would flux through defined ion channels. We modified the ion channel parameters to reflect the conductance profiles and capacitances observed in representative type I and type II SGNs to create a model type I and a model type II SGN. Additionally, sodium, leak, and HCN channel conductance densities were taken from previous models (Rothman and Manis, 2003; Hight and Kalluri, 2016; Ventura and Kalluri, 2019) and were equal in both of the simulated neurons. Four parameters were systematically varied across SGN subtypes: *g_KL_*, *g_KH_*, *g_KA_*, and membrane capacitance. These values were based on experimentally derived capacitances and conductance densities from representative cells (see Table 2 for full parameters).

Voltage-clamp simulations reproduced the current profiles observed in real neurons. Decomposing the total current into its three component potassium currents revealed that model type I SGNs relied primarily on *I_KL_* and *I_KH_*, with minimal contribution from *I_KA_* (Figure 9A-B). In contrast, the type II SGN model had reduced *I_KL_* and *I_KH_*, and prominent but transient I*_KA_*, which quickly inactivated.

**Figure 9:**
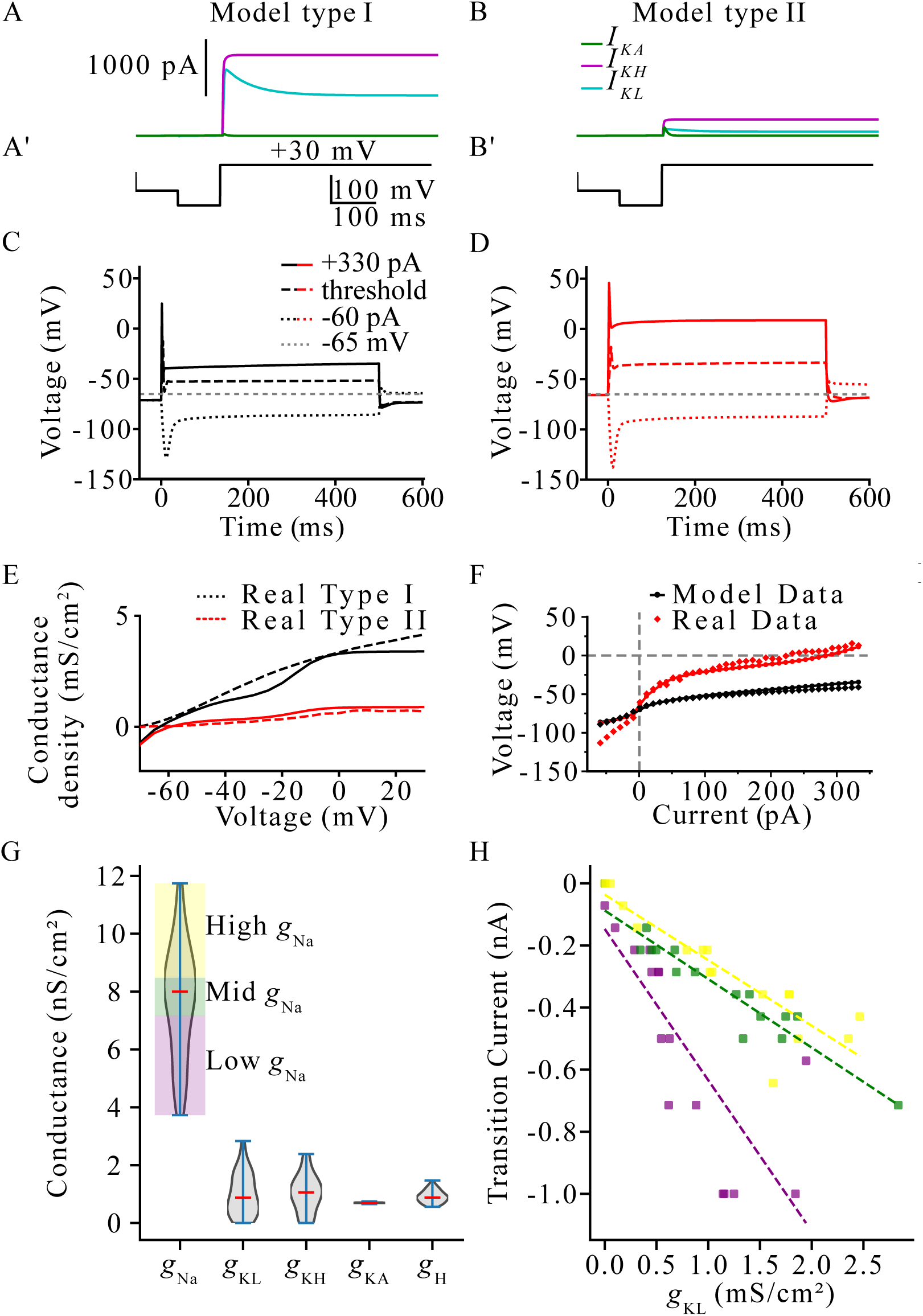
A conductance-based model replicates SGN behavior. (A–B) Simulated voltage-clamp traces for a model type I SGN (A) and type II SGN(B); green lines show example traces of the inactivating A-type K^+^ current (*g_KA_*), magenta line show high-threshold K^+^ curbrent (*g_KH_*), blue lines show low-threshold K^+^ current (*g_KL_*). (A’–B’) Voltage step protocols. (C–D) Current-clamp simulations with 500 ms steps; solid line equals maximum current (+330 pA), long dashed line equals threshold current, short dashed line equals minimum current (−60 pA), and gray dashed = -65 mV reference. (E) Total conductance scaled by size vs voltage; solid = simulated data, dashed = real data. (F) Steady-state voltage at 450 ms vs. current injection; diamonds = real data, circles with line = model data. Gray dashed lines = 0 values. (G) Range of independently drawn conductances for 50 simulated neurons. (H) Hyperpolarizing current required for anode break spike vs. *g_KL_*; fit lines for high/mid/low *g_Na_*: yellow, green, and magenta, respectively.

Having established that the model accurately captures the outward current profiles observed in real SGNs under voltage clamp, we next tested whether it could also reproduce key features of neuronal excitability under current-clamp conditions (Figure 9C–D). At rest, the type II SGN model settled at a more depolarized membrane potential than the type I SGN model, consistent with experimental findings. Both model neurons fired action potentials in response to extended square current pulses, but the type II SGN required less injected current (40 pA vs. 100 pA). The model type II SGN also showed larger voltage deflections for a given current, replicating the voltage–current relationship observed experimentally at steady state (diamonds, Figure 9F).

Altogether, the models faithfully reproduce the observed total conductance (Figure 9E) and current values (Figure 9F) of the real cells at each voltage level. However, the model type II SGN performed worst at estimating the cell’s voltage when hyperpolarizing current was injected. The more depolarized currents may explain why we did not observe any anode break spikes in the simulated current clamp experiments at least up to -60 pA. This may reflect differences in leak and or hyperpolarization activated currents that were not individually tuned.

To investigate anode-break spiking in more detail, we randomly generated 50 simulated neurons where each conductance was independently assigned from a normal distribution. This ensured that there was no confounding correlation between conductances. Figure 9G shows the distribution of assigned conductances in the simulated neurons. Next, each cell was presented progressively hyperpolarizing levels of current to determine the threshold for anode-break spiking. We applied an ANCOVA model to test whether anode-break threshold current depends on low-voltage–gated potassium conductance (*g_KL_*) and sodium conductance (*g_Na_*), with modeled neurons grouped into low-, medium-, and high-*g_Na_* density categories. The overall model was highly significant, with *g_KL_* and *g_Na_* together accounting for 75% of the variance in anode-break thresholds (F-statistic = 27.31, p-value = 1.84e-12). Anode-break thresholds decreased linearly with increasing *g_KL_* (p < 0.001), indicating that *g*_KL_ is a strong predictor of anode-break spiking. The slope of this relationship also depended significantly on the *g_Na_* group (p = 0.001): neurons with low *g_Na_* showed a steeper dependence on *g_KL_*, whereas neurons with high *g_Na_* exhibited a shallower slope. This interaction indicates that *g_Na_* moderates the influence of *g_KL_* on anode-break excitability (Figure 9H). Only cells with *g_KL_* less than 0.5 mS/cm^2^ produced an anode break spike at currents more positive than -200 pA. Overall, we conclude that despite having lower sodium conductance, the main driver of anode breaking spiking behavior in type II-S SGNs is the relative low density of low-voltage gated potassium conductance.

### Eliminating potassium channel inactivation permits repetitive firing in model type II SGNs

Having established that potassium conductance density accounts for many of the passive and single-spike properties of SGNs, we next asked whether these same factors could also explain differences in repetitive firing behavior between subtypes. To investigate this, we used the computational model described in Figure 9 to test a key functional distinction: the ability of type I SGNs, but not type II SGNs, to sustain high-frequency firing. We first set to find the maximum rate of spiking for the model neurons to assess if the model could recapitulate the firing patterns seen *in vitro*.

To determine the maximum rate of firing we systematically increased the time interval in 1 ms increments between two suprathreshold (450 pA) 3ms long current impulses and assessed the minimum interval where the neuron could fire a second spike. The model type I SGN fires a second action potential 7 ms after the initial suprathreshold current step, resulting in a maximum instantaneous firing rate of ∼140 Hz (Figure 10A). The model type II SGN on the other hand, only produced a second spike after 14 ms had elapsed for an instantaneous firing rate of ∼70 Hz (Figure 10B). These values align well with the empirical data where all cells could maintain a firing rate of 20 Hz but only type I SGNs could fire to every current impulse at 100 Hz.

**Figure 10:**
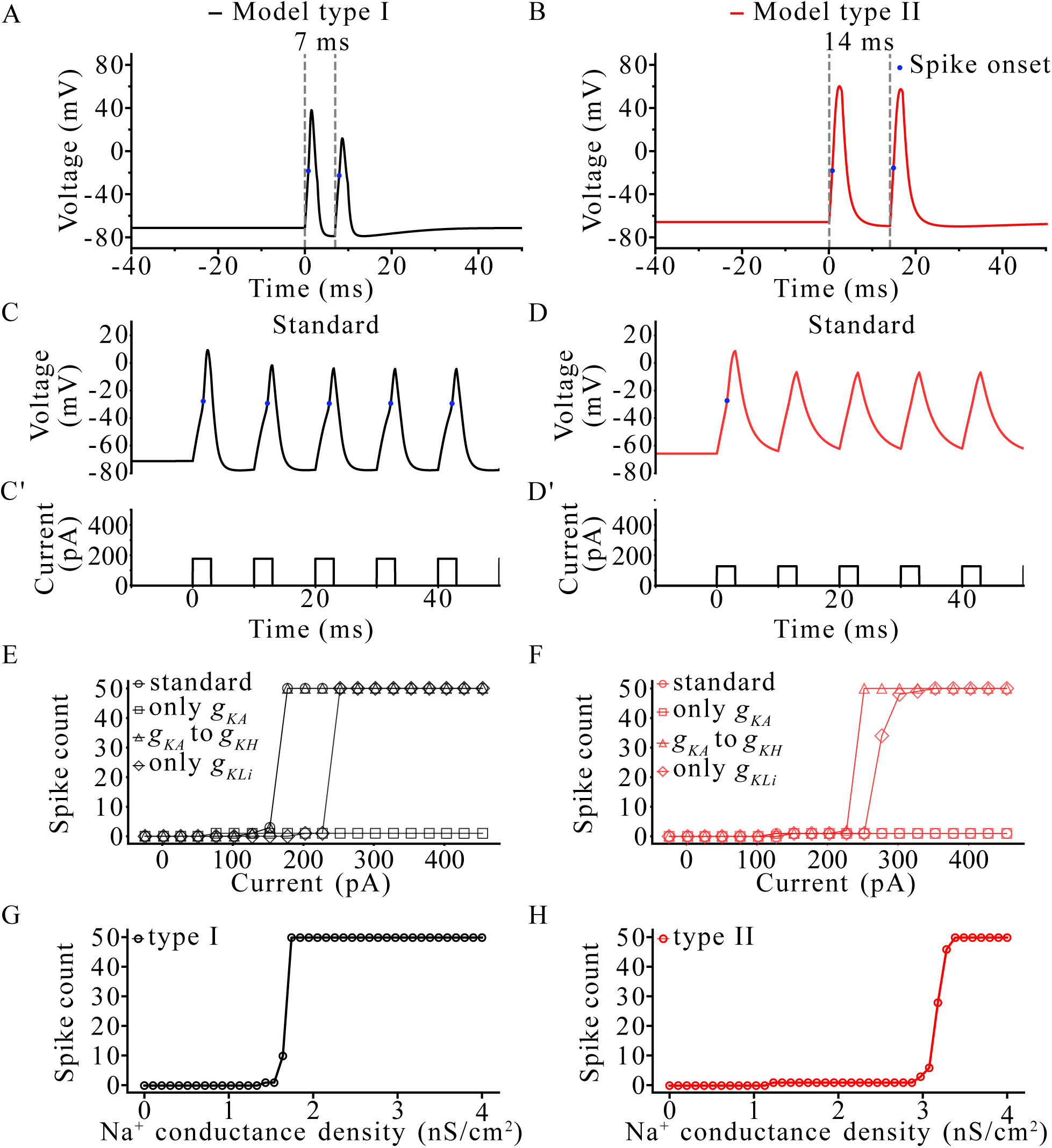
Modifying K^+^ and Na^+^ channels enables repetitive firing in type II SGNs. (A–B) Minimum inter-spike interval to produce second action potential; dashed = stimulus onset; blue dot = detected onset of action potential. (C–D) First five spikes of 100 Hz train at minimal current for the model type I SGN (C) and model type II SGN (D); (C’–D’) stimulus protocol. (E–F) Total spikes over 50 pulses for standard and K+ modification models for the model type I SGN (E) and model type II SGN (F): Circles = results for the standard model, squares = results for models where all K^+^ conductances are converted to the inactivating A-type K^+^ conductance (only *g_KA_*), triangles = results for models where the inactivating A-type K^+^ conductance is converted to the high-threshold K^+^ conductance (*g_KA_* to *g_KH_*), and diamonds = results for models where all K^+^ conductances are converted to a fully inactivating form of low-threshold K^+^ conductance (only *g_KLi_*). (G–H) Total spikes vs. Na^+^ channel conductance density for the model type I SGN (G) and model type II SGN (H).

Next, we simulated fifty 3ms current pulses delivered at 100 Hz, with amplitudes ranging from – 25 to +450 pA in 25 pA steps, the same stimulus protocol used in our experiments (Figure 8). In this simulation, the model type I SGN fired reliably to every suprathreshold pulse (Figure 10C), consistent with recorded neurons. In contrast, the model type II SGN generated only up to a single action potential regardless of current intensity (Figure 10D). Action potentials were detected using a custom algorithm that identified events where the second derivative of voltage (d²V/dt²) exceeded 45 mV/ms², indicating a rapid upswing consistent with sodium channel activation. Thus, the models faithfully reproduced the spiking phenotypes observed experimentally.

We then tested whether potassium current composition influenced the ability to fire repetitively. To isolate the effect of potassium current kinetics, we created three modified versions of both model neurons (Table 3). In the first variant, we converted all of the high-threshold (*g_KH_*) and low-threshold potassium (*g_KL_*) conductances into the inactivating potassium conductance (*g_KA_*). This allowed us to test whether type I SGNs with their larger total potassium conductance could overcome the inactivation of *g_KA_* to perpetuate their high frequency firing. In the second variant we shifted all *g_KA_* conductance into the non-inactivating *g_KH_* conductance. This model tested whether reducing the amount of potassium channel inactivation in type II SGNs could rescue their poor high frequency spike following. Lastly, we converted the *g_KA_* and *g_KH_* conductances into *g_KL_* and modified the *g_KL_* channel parameters such that the channel fully inactivated (*g_KLi_*). The final model tests whether the slower potassium channel inactivation of *g_KL_* can sustain high-frequency firing where *g_KA_* could not despite both fully inactivating to sustained activation.

**Table 3:**
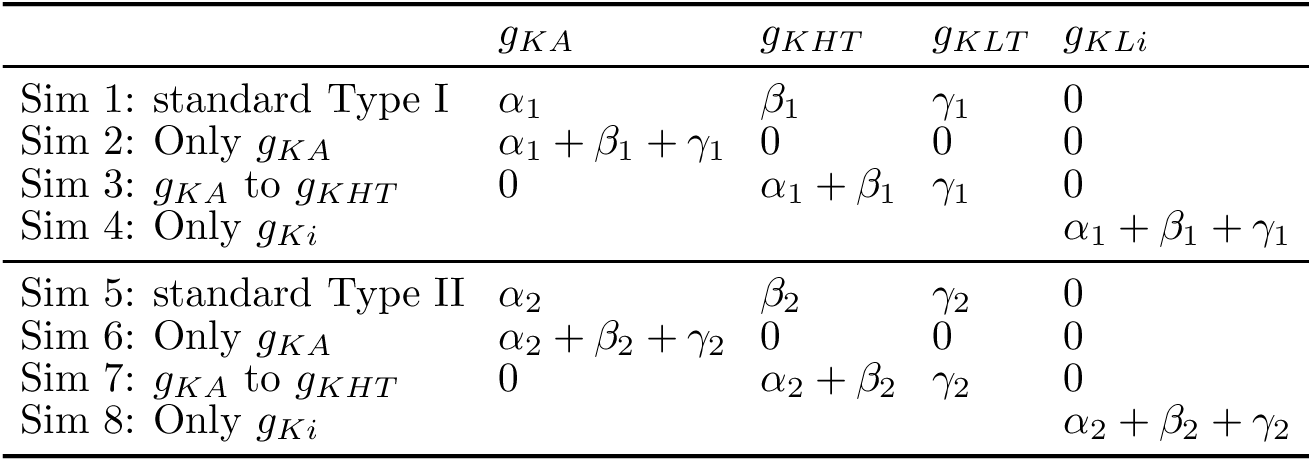
K conductance values in Type I and Type II simulations of Figure 10. For Type I: α_1_ = 0.65 nS/pF, β_1_ = 2.09 nS/pF, γ_1_ = 2.09 nS/pF. For Type II: α_2_ = 0.69 nS/pF, β_2_ = 0.5 nS/pF, γ_2_ = 0.25 nS/pF.

| | $g_{KA}$ | $g_{KHT}$ | $g_{KLT}$ | $g_{KLi}$ |
| --- | --- | --- | --- | --- |
| Sim 1: standard Type I | $\alpha_1$ | $\beta_1$ | $\gamma_1$ | 0 |
| Sim 2: Only $g_{KA}$ | $\alpha_1 + \beta_1 + \gamma_1$ | 0 | 0 | 0 |
| Sim 3: $g_{KA}$ to $g_{KHT}$ | 0 | $\alpha_1 + \beta_1$ | $\gamma_1$ | 0 |
| Sim 4: Only $g_{Ki}$ | | | | $\alpha_1 + \beta_1 + \gamma_1$ |
| Sim 5: standard Type II | $\alpha_2$ | $\beta_2$ | $\gamma_2$ | 0 |
| Sim 6: Only $g_{KA}$ | $\alpha_2 + \beta_2 + \gamma_2$ | 0 | 0 | 0 |
| Sim 7: $g_{KA}$ to $g_{KHT}$ | 0 | $\alpha_2 + \beta_2$ | $\gamma_2$ | 0 |
| Sim 8: Only $g_{Ki}$ | | | | $\alpha_2 + \beta_2 + \gamma_2$ |

In the model type I SGN, redistributing potassium conductance to only contain *g_KA_* prevented the cell from firing after the initial spike just like the model type II SGN, suggesting that the overall potassium conductance density was insufficient to sustain firing regardless of inactivation kinetics. Furthermore, as expected, eliminating *g_KA_* and adding to *g_KH_* had no change as compared to the standard model. Finally, type I SGNs fired 50 out of 50 spikes even when the only potassium conductance present was the fully inactivating *g_KLi_* but with an increase in the threshold to first fire (Figure 10E).

For the model type II SGN, converting all potassium conductance into *g_KA_* had no additional effect on the spiking behavior. However, converting all *g_KA_* into *g_KH_* enabled firing to each suprathreshold pulse in the 100 Hz train. Furthermore, when the only potassium conductance is the fully inactivating *g_KLi_* type II SGNs could fire to 100 Hz (Figure 10F). This result suggests that the rapid inactivation of potassium currents in type II SGNs, particularly in type II-S SGNs, impairs their ability to repolarize the membrane potential efficiently and thus cannot reset sodium channels for subsequent spikes.

Lastly, we set to determine whether the concentration of sodium conductance can influence the spiking rate of each model neuron. For the model type I SGNs, the minimum amount of *g_Na_* required to fire any action potentials at 450 pA was ∼1.4 mS/cm^2^ and to fire all 50 action potentials was ∼1.7 mS/cm^2^ (Figure 10G). Type II SGNs despite having a lower threshold for a single spike (∼1.2 mS/cm^2^) required much more sodium conductance to overcome the effects of their potassium conductances (∼3.4 mS/cm^2^) (Figure 10H).

Together, these simulations demonstrate that the inability of type II SGNs to fire repetitively arises both from differences in availability of sodium currents and from a lack of sustained potassium currents. Specifically, for neurons like type II-S SGNs, this combination of low sodium conductance and rapidly inactivating potassium conductance limits their capacity to follow high-frequency stimulation. Furthermore, the differences in the contour of the sodium conductance plots in Figure 10 G-H and imply an interaction between sodium and potassium conductances where beven more sodium conductance is needed to overcome the poor repolarization ability of type II SGNs.

## Discussion

### Genetic tools improve the targeting of type II SGNs

Efforts to define type II SGN physiology have long been limited by the difficulty of reliably identifying these rare neurons. Early studies relied on immunolabeling or sparse anatomical markers, leading to small samples and uncertain cell identity. Reid et al. (2004) used peripherin to target type II SGNs but recorded only eight neurons, likely including immature type I SGNs. Markowitz et al. (2020) labeled single fibers in whole mounts and also obtained eight neurons across a wider developmental range. Jagger and Housley (2003) used a demanding slice preparation but definitively identified only eight type II SGNs, plus twenty putative cells. Comparisons across studies are further complicated by differences in culture duration, enzymatic treatments, species, and developmental stage.

Type II SGN–biased Cre reporter lines address many of these limitations by providing specific, scalable targeting. These tools enabled us to record from a sufficiently large population to distinguish type I and type II SGNs and to resolve electrophysiological subgroups within each. Our findings thus provide a more comprehensive view of type II SGN biophysical diversity and establish a modeling framework for future work.

### Biophysical differences between type I and type II SGNs

To compare type II and type I SGNs, we used two previously characterized type II SGN–selective fluorescent reporter lines (SERT^Cre^; Ai14 and Tac1^Cre^; Ai14; Vyas et al., 2019; Nowak et al., 2021). Both lines were strongly biased toward type II SGNs, as confirmed by Tuj1 immunostaining (85% and 46% labeled, respectively, compared with 5% by chance). Electrophysiological recordings revealed two broad biophysical groups corresponding to these labeling proportions: nearly 80% of SERT^Cre^; Ai14 neurons exhibited a type II SGN–like profile, whereas about half of Tac1^Cre^; Ai14 neurons did.

Type II SGNs had smaller outward potassium currents, a substantial portion of which inactivated over time, and reduced inward sodium currents compared to type I SGNs. These features limit repetitive firing and slow action potential repolarization, contributing to the relative quiescence of type II SGNs *in vivo* (Robertson, 1984; Brown, 1994; Flores et al., 2015; Weisz et al., 2021).

### Biophysical diversity within type I and type II SGN populations

Biophysical diversity among type I SGNs is well documented and correlates with synaptic position along the inner hair cell (Markowitz and Kalluri, 2020) and with tonotopic location (Adamson et al., 2002). Variability in potassium conductances produces differences in firing behavior, including spike latency and threshold. The gradients in outward conductance, spike latency, and current threshold in our recordings align with the modiolar–pillar patterns described previously, indicating that our dataset captures the full spectrum of type I SGN subtypes.

Transcriptomic gradients in type II SGNs suggest potential subtypes related to tonotopic position (Vyas et al., 2019), including apical–basal biases in *Tyrosine Hydroxylase* and *Calca* expression (Wu et al., 2018). Although the SERT^Cre^; Ai14 line is apically biased, we recorded SERT+ neurons from both cochlear turns and complemented these data with Tac1^Cre^; Ai14 mice, which label type II SGNs uniformly (Nowak et al., 2021). Across all recordings, we found no significant biophysical differences between apical and basal neurons, indicating that transcriptomic gradients do not translate to electrophysiological gradients.

### Cell size as a marker and correlate of diversity within type II SGNs

Type II SGNs are often described as smaller than type I SGNs, but our data and previous anatomical work show that they span a broader size range (67–176 µm² vs. 102–149 µm²). Some type II SGN somata were comparable to or larger than type I SGNs, indicating that cell size alone cannot reliably distinguish the two types.

Within the type II SGN population, soma area and capacitance were the strongest correlates of biophysical diversity. Small type II-S SGNs (107 µm²) showed pronounced potassium-channel inactivation and frequent anode-break spiking. Medium-sized type II-M SGNs (130 µm²) resembled type I SGNs in size but differed in potassium current magnitude and inactivation. Jagger and Housley (2003) noted smaller (∼101 µm²) and larger (∼148 µm²) type II SGNs but did not link size to function. Our data provide the first functional characterization of these subgroups.

We also identified three exceptionally large SGNs, one reaching 216 µm², which generated full action potentials and anode-break spikes with the largest observed sodium and potassium conductances. Rare large type II SGNs have been reported (Berglund and Ryugo, 1987; Mo and Davis, 1997; Pearson et al., 2025) and likely represent a distinct II-L SGN subtype. Overall, our findings support three size-defined type II SGN subtypes: II-S, II-M, and II-L, each with distinct conductances and firing behaviors.

### Identity of the inactivating potassium current (*I_KA_*)

A prominent feature distinguishing type II-M and particularly type II-S SGNs was an inactivating outward current. Jagger and Housley (2003) described such a current and noted its sensitivity to 4-aminopyridine (4-AP), an antagonist of voltage-gated potassium channels (Grissmer et al., 1994; Zemel et al., 2018). A likely molecular candidate is *KCNC4* (Kv3.4), an inactivating potassium channel selectively enriched in type II SGNs (Shrestha et al., 2018). Immunohistochemistry confirms preferential Kv3.4 expression in type II SGNs (Kim et al., 2020). Kv3.4 channels are 4-AP–sensitive and inactivate near –25 mV (Ritter et al., 2012; Zemel et al., 2018), consistent with the pharmacological profile of the inactivating current observed. Thus, Kv3.4 is the most plausible mediator of the *I_KA_* that constitutes up to 40% of outward conductance in some type II SGNs.

### Parallels between spiral ganglion and dorsal root ganglion neurons

Dorsal root ganglion (DRG) neurons detect multiple somatosensory modalities (Abraira and Ginty, 2013). Nociceptors (small, unmyelinated, damage-responsive neurons) often express neuropeptides such as substance P (*Tac1*) and CGRP (*Calca*) (Basbaum et al., 2009; Usoskin et al., 2015). Type II SGNs share these features: they are small, unmyelinated, respond to damage-induced ATP release (Liu et al., 2015; Nowak et al., 2021), and are selectively labeled by *Calca*-and *Tac1*-based reporter lines (Wu et al., 2018; Nowak et al., 2021; Sanders and Kelley, 2022).

Kv3.4 channels, also present in nociceptive DRG neurons (Ritter et al., 2012), strengthen these parallels. After nerve injury, PKC-dependent phosphorylation suppresses Kv3.4 inactivation, converting phasic to sustained firing (Ritter et al., 2015). Type II SGNs likewise become hyperactive after traumatic noise exposure (Nowak et al., 2021), perhaps via similar modulation. Our modeling shows that removing *I_KA_* inactivation converts type II SGNs from single-spike responders to neurons firing >100 spikes/s, implicating Kv3.4 and PKC signaling as potential mediators of noise-induced hyperexcitability.

We also observed differences in sodium currents between SGN types. Type II SGNs had smaller peak currents and trended toward more depolarized half-activation voltages. Although series-resistance artifacts warrant caution, these findings raise questions about Nav isoform composition. Most work focuses on Nav1.6 and β subunits (Hossain et al., 2005; Browne et al., 2017) but transcriptomic data show low-level expression of additional isoforms, including Nav1.1, Nav1.2, and Nav1.7, although the small number of captured type II SGNs prevents subtype-specific comparisons (Petitpré et al., 2018; Shrestha et al., 2018; Sun et al., 2018). Given parallels with DRG neurons, where Nav isoforms shift dynamically during injury or pain (Waxman et al., 1999; Cummins et al., 2007), defining sodium-channel composition in SGN subtypes is an important goal for future studies.

### Possible stimuli to drive type II SGN excitability

Most type II SGNs did not spike at rest, consistent with sodium-channel inactivation at their relatively depolarized membrane potentials. Although spike initiation normally occurs in the unmyelinated dendrites, somatic recordings nevertheless reflect functional consequences of channel availability. Many type II SGNs fired following transient hyperpolarization, suggesting that inhibitory inputs could prime them for activity. Type II SGNs express GABA_A_ receptors and may receive medial olivocochlear efferent input (Bachman et al., 2025), which could enable rebound spiking if these receptors become hyperpolarizing in adults, as occurs in many other neurons (Cherubini et al., 1991; Ouardouz and Sastry, 2005).

In addition, type II SGNs have large membrane time constants due to either high input resistance (II-S, II-M) or high capacitance (II-L), making them well suited to integrate slow or spatially distributed inputs, consistent with their broad outer hair cell innervation (Weisz et al., 2009, 2012). This property may be particularly relevant for detecting sustained ATP release from damaged outer hair cells, a known stimulus for type II SGNs (Liu et al., 2015; Nowak et al., 2021). Together, these features suggest that type II SGNs are tuned to detect weak, sustained cochlear signals, such as damage-related cues, rather than the rapid, transient inputs that drive type I SGNs (Siegel, 1992; Rutherford et al., 2012).

### Toward a functional characterization of type II SGNs

The wide variation in ionic conductances, soma size, and firing patterns among type II SGNs suggests multiple functional subtypes, paralleling diversity observed in DRG neurons. Future studies combining genetic labeling, *in vivo* recording, and behaviorally relevant stimulation will be essential for determining what each subtype detects and how these signals influence auditory perception. Understanding this spectrum of properties will provide a foundation for linking molecular identity with function and for exploring how type II SGNs contribute to normal and pathological auditory processing.

## Acknowledgements

This work was supported by NIH NIDCD Grants R01 DC015512 and T32 DC009975, and the Caruso Department of Otolaryngology at the University of Southern California. We thank Dr. Daniel Bronson, Katherine Regalado and members of the Kalluri PIELab @ USC, Department of Otolaryngology, and the Hearing and Communications Neuroscience Training Program for discussion. We thank Juan Llamas and Dr. Ksenia Gnedeva for teaching us the intracerebroventricular approach for injecting viral (AAV) vectors.

